# Molecular Dynamics in the Ventral Tegmental Area during Chronic Pain-Induced Negative Affect

**DOI:** 10.1101/2025.07.01.662604

**Authors:** Cody C. Diezel, Lisa A. Majuta, Erfan Bahramnejad, Kelly L. Karlage, Jennifer L. Partin, Saniya M.D. Barbour, Ingrid L. Peterson, Matthew Flowers, Sophia T. von Hippel, Riley Haveman, Ethan Villarroel, Isabella Villarroel, Paul R. Langlais, Tally M. Largent-Milnes, Todd W. Vanderah, Arthur C. Riegel

**Author notes:** **To whom correspondence should be addressed:** Arthur C. Riegel, Department of Pharmacology, University of Arizona, 1501 N. Campbell Ave, Tucson AZ 85724, United States.

## Abstract

Chronic pain frequently coexists with negative affect, with about 60% of patients suffering from both. This dual condition complicates treatment and exacerbates both disorders, highlighting the urgent need for innovative therapeutic strategies. Chronic pain negative affect (CPNA) involves complex neurobiological changes, including increased hyperexcitability of the ventral tegmental area (VTA), a critical region involved in reward, mood, and pain processing. To elucidate CPNA’s underlying mechanisms, we employed a multidisciplinary approach using immunohistochemistry, lipidomic analysis, and proteomic screening to investigate VTA molecular alterations in mice subjected to partial sciatic nerve ligation (pSNL) at one and four weeks post-injury. Our results revealed a significant, sex-dependent increase in Kv7.2 channel expression in dopamine neurons, alongside a notable reduction in endocannabinoid 2-arachidonoylglycerol (2-AG) levels, which plays a vital role in mood regulation. This neurochemical shift associated with an increase in negative affect-like behaviors, as determined by the forced swim test. Furthermore, pharmacological intervention utilizing either exogenous 2-AG or retigabine, a Kv7 channel opener, effectively alleviated pain-related negative affect symptoms. Proteomic profiling further uncovered alterations within the CaMKK2 pathway, involving crucial proteins such as PLCγ2, AMPKγ2, and AMPKβ1, and CaMK1α, with changes in abundance and phosphorylation activity that could be reversed with the next-generation CaMK1α antagonist CS640. This research provides the first comprehensive analyses of VTA adaptations linked to CPNA, yielding significant insights into molecular changes impacting VTA neuronal integrity and signaling throughout CPNA progression.

## Introduction

Chronic pain, persistent discomfort lasting three months or longer [1], is a complex phenomenon deeply intertwined with psychological conditions, particularly negative affect. Nearly 60% of chronic pain patients also experience clinically significant negative affect symptoms, highlighting a profound intersection between these debilitating conditions [2, 3]. This bidirectional association—where chronic pain exacerbates negative affect and negative affect heightens pain perception—creates a vicious cycle, complicating treatment and significantly reducing quality of life [2, 4, 5]. Understanding this intricate relationship is crucial for developing effective treatment strategies addressing both chronic pain and comorbid negative affect.

The ventral tegmental area (VTA), a key brain region for motivation and reinforcement, has been identified in recent chronic pain negative affect (CPNA) mechanism investigations [4, 5]. The VTA is pivotal not only in motivational behaviors but also in pain modulation and mood regulation [6]. Alterations in VTA dopaminergic neuron excitability occur under pain and negative affect conditions. However, mixed findings exist regarding VTA hyperexcitability [7–11] or hypoexcitability [11–18]. For instance, a human fMRI study indicated heightened VTA-medial prefrontal cortex (mPFC) connectivity in major depressive disorder (MDD) subjects, suggesting increased activity in pathways where pain may modulate VTA function and influence mood [7]. Preclinical models support these findings, showing VTA hyperactivity in rodents exhibiting negative affect-like behaviors weeks after neuropathic injury [8, 9]. Retigabine, a positive allosteric modulator (PAM) of Kv7/KCNQ channels that mediate the M-current, effectively reduces VTA hyperactivity and improves negative affect-like behaviors in these models [8, 9]. These observations underscore the pivotal role of VTA neuronal excitability changes in developing negative affect within CPNA models. Kv7 channels, comprising four brain-expressed subunits (Kv7.2-Kv7.5), significantly influence neuronal excitability and behavior [19]. However, comprehensive evaluations of all Kv7 subunits’ expression alterations in the VTA within the context of chronic pain and negative affect remain limited.

Recent research explores the intricate relationships among Ca2+/calmodulin-dependent protein kinase kinase 2 (CaMKK2), calcium/calmodulin-dependent protein kinase 1α (CaMK1α), and AMP-activated protein kinase (AMPK) in chronic pain [20–24]. CaMKK2, a critical serine/threonine protein kinase, is involved in calcium signaling, influencing neuronal growth, synaptic plasticity [20], and mood regulation [20, 25]. Mood-stabilizing medications enhance CaMKK2 activity and expression [25]. Activated by calcium and calmodulin, CaMKK2 initiates CaMK1α and AMPK activation, which have opposing morphogenic roles [22, 26–30]. CaMK1α supports synapse formation by phosphorylating postsynaptic density proteins, including NMDA receptors critical for long-term potentiation [20, 21]. Next-generation CaMK1 inhibitors, such as CS640, are potential tools to investigate neuropsychiatric illness [25, 31]. Conversely, AMPK reduces energy metabolism and neuronal activity while enhancing autophagy by phosphorylating microtubule-associated proteins, opposing CaMK1α effects [26]. Thus, CaMKK2 signaling dysregulation in chronic pain and negative affect may influence VTA neuronal morphogenesis and excitability, impacting negative affect development. The interplay between CaMKK2, CaMK1α, AMPK, and Kv7 channels in the VTA may yield insights into the molecular mechanisms underlying persistent pain’s emotional aspects.

The endogenous cannabinoid system, particularly 2-arachidonoylglycerol (2-AG), is an intriguing element in this complex interplay. Acute pain increases 2-AG levels [32–34], yet reduced levels are seen in individuals with negative affect [35–37], suggesting eCB signaling disruption in chronic pain and mood disorders. In non-neuronal cells, 2-AG activates the CaMKK-AMPK pathway to control phosphorylation and actin polymerization [38, 39]. In male hypothalamic neurons, 2-AG synthesis links to increased intracellular calcium and CaMKK-AMPK signaling activation [40]. Other studies identified endocannabinoids, including 2-AG and phytocannabinoids like CBD, as potential PAMs for Kv7 channels [41–43], potentially linking endocannabinoid mechanisms to VTA neuronal excitability modulation. However, how chronic pain alters VTA activity and subsequently develops negative affect symptoms through these mechanisms remains unclear, necessitating further research.

To bridge existing knowledge gaps, we employed a partial sciatic nerve ligation (pSNL) model to induce neuropathic pain in mice, enabling comprehensive exploration of VTA alterations at one and four weeks post-surgery. We investigated molecular alterations using immunohistochemistry (IHC), lipidomic analysis, and proteomic/phosphoproteomic strategies to probe protein and endocannabinoid levels. Behavioral impacts were assessed using von Frey (vF) tests for paw hypersensitivity and the forced swim test (FST) for negative affect-like behaviors. Our proteomic examinations also evaluated early protein changes shortly after pSNL and at the four-week interval, offering insights into how these alterations influence CPNA development and disrupt VTA functioning. This research aimed to illuminate VTA molecular changes in CPNA, unraveling mechanisms intertwining chronic pain and negative affect symptoms, thereby establishing a framework for future investigations.

## Methods

For more detailed methods and vendor information, see **Supplement Material 1.**

### Animals

A total of 154 (138 male, 16 female) C57BL/6J mice (Jackson Labs), aged 7-8 weeks, were housed three to five per cage in standard cages with ad libitum access to food and water, under a 12-hour light-dark cycle in a climate-controlled room. All procedures were approved by the University of Arizona Animal Care Use Committee and complied with NIH guidelines for laboratory animal care.

### Partial Sciatic Nerve Ligation

All 154 mice underwent surgery [44], with 80 receiving pSNL and 74 undergoing sham operations, following published procedures. Mice were anesthetized and administered gentamicin at a dose of 8 mg/kg (IP).

### Mechanical Withdrawal Threshold

Tactile allodynia was evaluated at baseline and on days 5 and 28 post-surgery by measuring the paw withdrawal threshold using calibrated von Frey filaments on the nerve injury side, following published procedures [45]. Tactile allodynia was defined as a significant drop in the withdrawal threshold compared to baseline [45].

### Forced Swim Test (FST)

Despair-like behavior was assessed 28 to 35 days post-surgery in male mice by measuring immobility time in a water-filled cylindrical container (25LJC) for up to 5 minutes. Total immobility time throughout the duration of the test was used to evaluate despair-like behavior. One cohort received 2-AG (10 mg/kg, IP), prepared in a 1:1:8 DMSO:Tween 80:saline solution, 32 days post-surgery, and RTG (10 mg/kg, IP), prepared in 0.9% saline, 35 days post-surgery, administered 15 minutes before the FST. Another cohort received two doses of 2-AG (10 mg/kg, IP) 31 days post-surgery, both 15 and 60 minutes before the FST.

### Immunohistochemistry (IHC)

VTA samples were processed for IHC similar to published procedures [46]. Samples were placed in primary antibody solutions: anti-KCNQ2 (1:800), anti-KCNQ3 (1:800), anti-KCNQ4 (1:400), anti-KCNQ5 (1:200), anti-CaMK1α (1:500), and anti-tyrosine hydroxylase (TH) (1:800). After primary incubation, tissues underwent PBS washes and were incubated with secondary antibodies: anti-Guinea Pig Cy3 (1:600) and anti-Rabbit AF488 (1:400). DAPI was used for counterstaining, and slides were mounted with Prolong Glass.

### Imaging and Analysis

Photomicrographs of tissue samples were taken using an Olympus FluoView FV1200 confocal microscope at 20x magnification. The VTA was identified via coronal section schematic overlays from the mouse brain atlas (Franklin & Paxinos, 3rd Edition, 2007). Analyses were conducted with FIJI/ImageJ software. Quantifiable values for TH+, Kv7+, TH+/Kv7+, CaMK1α+, and TH+/CaMK1α+ were obtained from 2-3 VTA slices contralateral to the surgical site for each mouse.

### Quantification of 2-AG and AEA by LC-MS

Pooled VTA samples (n=4−5 bilateral punches/sample) underwent organic solvent extraction for LC-MS analysis following established methodologies. Mixed internal standards were created via serial dilution of d4-AEA and d5-2-AG in acetonitrile. Analysis of 2-AG and AEA was performed using an Agilent 6495C triple quadrupole mass spectrometer linked to a 1 290 Infinity II UPLC system. All samples were run in triplicate.

### Label-Free Quantitative Proteomics

Pooled VTA samples (2-3 bilateral punches/sample) from 1 and 4 weeks post-surgery were processed for all proteome-wide experiments following previously published procedures [47–49]. 50 µg of VTA lysate was separated by SDS-PAGE, purified, and prepared for MS analysis as previously described [47, 48]. HPLC-ESI-MS/MS and label-free quantitative proteomics were conducted in accordance with the aforementioned procedures [48]. Unbiased hierarchical clustering analysis (heat map) and principal component analysis (PCA) were conducted using Perseus. Gene ontology (GO) and Reactome pathway enrichment analyses were completed with the DAVID tool. For synapse-specific subcellular localization analysis, SynGO was utilized. Phosphorylation sequence enrichment analysis was performed using the iceLogo generation tool. Volcano plots and scatter plots were created in GraphPad Prism.

### Phosphoproteomics

To evaluate differences in protein phosphorylation abundance with pSNL and CS640 treatment, mice were anesthetized and administered a 5 µL intracerebroventricular injection of CS640 (2 µg/µl) or vehicle (1:1:8 DMSO:Tween 80:saline solution). The researchers that administered the CS640 and vehicle were blinded to group assignments. Bilateral VTAs were collected 30 minutes later. 0.2-0.4 mg of VTA protein lysate per sample underwent in-solution tryptic digestion and phosphopeptide enrichment using metal oxide affinity chromatography per the manufacturer’s protocol, similar to as previously described [50, 51]. The dried peptides were resuspended in 20 µl of 0.1% FA (v/v), and peptide concentration was determined with the Pierce Quantitative Colorimetric Peptide Assay Kit (ThermoFisher Scientific). 350 ng of the final sample was then analyzed by mass spectrometry.

### Statistical Analysis

Statistical analyses were conducted using GraphPad Prism 10.4.2, employing two- and three-way (mixed-effects (ME) as necessary) ANOVAs, with Tukey’s or Fisher’s LSD post hoc tests for group comparisons and unpaired t-tests where appropriate. All GraphPad Prism statistics are detailed in **Supplementary Table 1**.

## Results

### Neuropathic Pain Lowers VTA Levels of 2-AG at Week 4

Neuralgia has been linked to VTA dopaminergic neuron hyperexcitability and negative affect-like behaviors [9, 10]. To assess whether a different neuropathic pain model produces similar effects, we performed pSNL and sham surgeries on mice. Mechanical sensitivity and immobility in the forced swim test (FST) were assessed prior to and four weeks post-surgery (**Fig. 1A**). VTA tissue was collected for lipidomic profiling, IHC, and proteomic analysis (**Fig. 1A,B**).

**Fig. 1.**
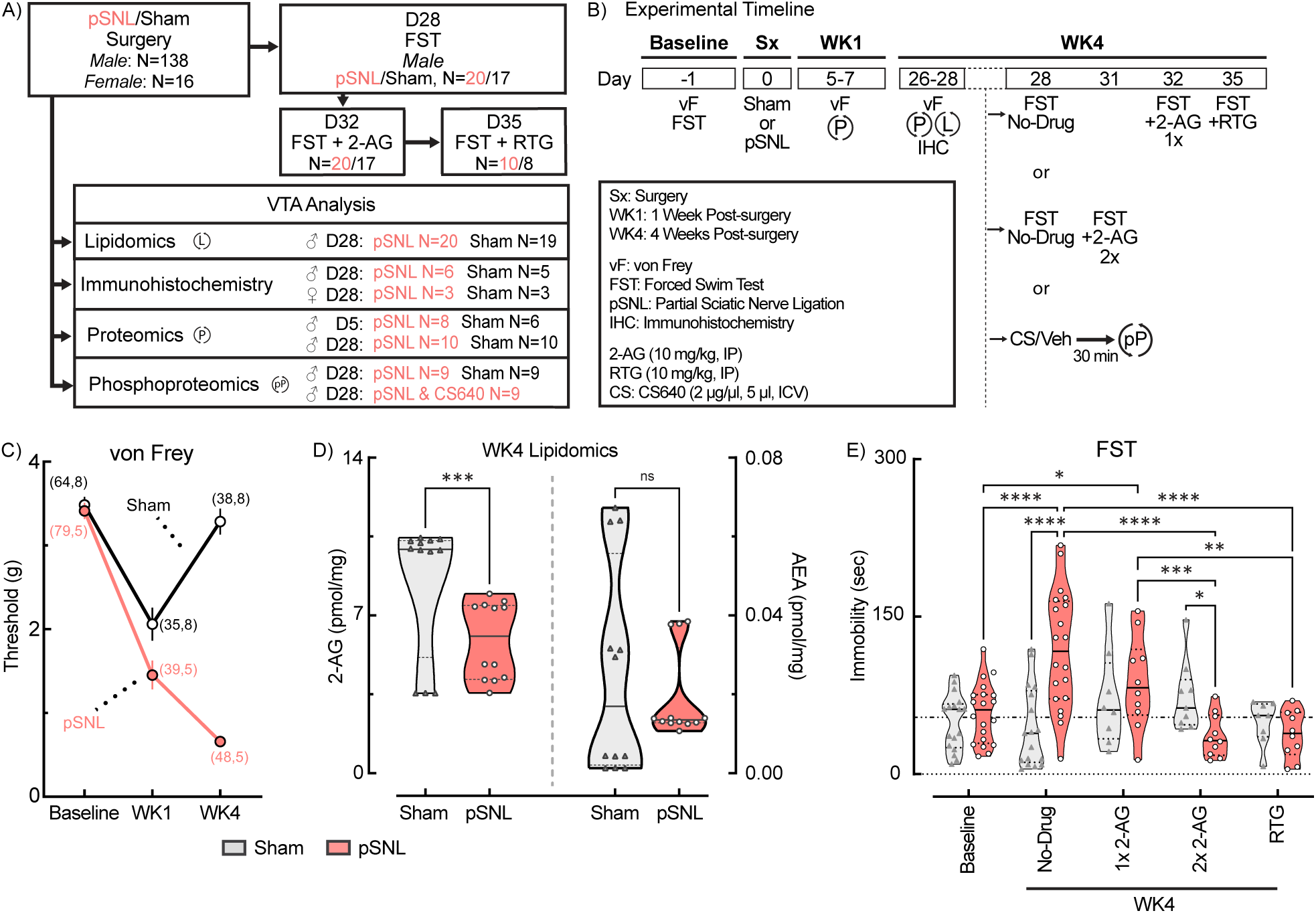
WK4 pSNL Alters VTA 2-AG Levels and Negative Affect-Like Behavior: Mitigation by 2-AG and Retigabine. **a, b** Schematic representation of group allocations and experimental timeline for cohorts of male and female mice subjected to either sham surgery or pSNL. Various VTA analyses, including IHC, proteomics, and lipidomics, were conducted at multiple timepoints. Behavioral assessments using von Frey (vF) tests and Forced Swim Tests (FST) were performed at baseline (pre-intervention), 5-7 days post-surgery (WK1), and 28 days post-operation (WK4). Some mice underwent additional FST evaluations in WK4 (day 28, 31, 32, or 35) either without drug (No-Drug) or 15 min after drug (single (1x) or double (2x) injection of 2-AG (10 mg/kg, IP), or single injection of retigabine (RTG; 10mg/kg, IP)). **c** Summary of paw withdrawal thresholds in male and female mice from sham and pSNL-exposed groups. Mechanical paw sensitivity analysis showed significant main effects of time, surgery type, and their interaction (2W ME ANOVA, F(2,151)=39.46, ***p<0.0001). Both groups exhibited hypersensitivity at WK1 compared to baseline (****p<0.0001). pSNL mice showed persistent hypersensitivity at WK4 (****p<0.0001), while sham mice resolved to baseline levels (p=0.2698). Sample sizes are indicated in parentheses (male, female). **d** Violin plots summarizing levels of 2-AG and anandamide (AEA) in male mice at WK4 post-surgery (2W RM ANOVA, interaction F(1,22)=6.810, *p=0.0160). pSNL mice exhibited lower levels of 2-AG compared to sham control mice (***p=0.0006), while AEA concentrations remained unchanged (p=0.9931) (see also **Supplementary Figure 1**). Samples were run in triplicate. **e** Results from FST measuring the duration of immobility in male mice at baseline and post-surgery at WK4. Immobility was assessed without drug or with 2-AG or RTG pretreatment (2W ME ANOVA, interaction F(4,84)=11.89, ****p<0.0001). pSNL mice exhibited significantly increased immobility at WK4 compared to baseline and sham controls (****p<0.0001). Both double-dose 2-AG and RTG significantly reduced FST immobility (both ****p<0.0001), while a single dose of 2-AG was less effective (****p=0.0006). All data are expressed as mean ± SEM. In all violin plots, solid lines depict median and dashed lines depict quartiles.

Mechanical paw sensitivity analysis revealed significant main effects of time, surgery type, and their interaction (2W ME ANOVA, all ****p<0.0001) (**Fig. 1C; Supplementary Figure 1**). pSNL mice exhibited persistent allodynia from WK1 to WK4 (WK1, ****p<0.0001; WK4, ****p<0.0001), while sham mice displayed transient allodynia that resolved by WK4 (WK1, ****p<0.0001; WK4, p=0.3197). Sex had no significant effect (3W ME ANOVA). These results affirm the pSNL model’s effectiveness for producing long-lasting allodynia [44].

While increases in 2-AG immediately following injury may predict future pain progression [32–34], subsequent reductions may indicate heightened risk for negative affect [33]. Our preliminary proteomic analysis at WK1 (**Supplementary Figure 2; Supplementary Table 2**) showed significant increases in eCB degradation enzymes FAAH (*p=0.0306) and ABHD6 (*p=0.0299) in pSNL mice, non-significant increases in synthetic eCB enzymes MGLL and DAGLA, decreases in KAP3 (*p=0.0292), and no changes in NAPEPLD or KCNQ2/Kv7.2 channels, possibly signaling a mild neuromodulatory imbalance in eCB signaling and glutamate plasticity during early WK1 pSNL [52, 53]. To examine eCB levels at WK4, LC/MS was utilized (**Fig. 1D**). Pooled bilateral VTA punches from four to five mice per group ensured adequate lipid extraction for eCB analysis. A 2W ANOVA revealed significant effects of surgery, eCB type, and their interaction on eCB concentrations (surgery: *p=0.0164; eCB type: ****p<0.0001; interaction: *p=0.0160). Fisher’s LSD post hoc analysis showed a significant reduction in 2-AG in pSNL mice (****p=0.0006), while anandamide (AEA) remained unchanged (p=0.9931). These findings are consistent with earlier reports that chronic pain alters eCB levels in various brain regions [53, 54].

### WK4 pSNL Increases Negative Affect Behavior

Low 2-AG levels have been linked to negative affect [35, 36, 55], and modeling studies suggest Kv7 channels activated by retigabine may also respond to 2-AG [41]. To determine if exogenous 2-AG or retigabine could reduce negative affect-like behaviors, we tested 37 mice in the FST before surgery (Day-1) and again four weeks post-surgery (**Fig. 1B**). During WK4 testing, immobility was assessed with pretreatments of a single dose of 2-AG (10 mg/kg, IP, 15 min prior), two doses of 2-AG (10 mg/kg, IP, 60 and 15 min prior), or retigabine (10 mg/kg, IP, 15 min prior). A 2W ME ANOVA of immobility demonstrated a significant interaction between surgery and drug treatment (drug: ****p<0.0001; surgery: p=0.3050; interaction: ****p<0.0001) (**Fig. 1E**). Tukey’s post hoc analysis revealed untreated pSNL mice exhibited increased immobility at WK4 compared to baseline and sham controls (****p<0.0001). Both double-dose 2-AG and retigabine significantly reversed chronic pain-induced negative affect (CPNA) immobility (****p<0.0001), while a single dose of 2-AG was less effective (***p=0.0006), indicating a possible dose-dependent effect. Sham-operated mice’s immobility was unchanged across all time points and treatments (p>0.05). These results expand upon previous retigabine studies [9, 10], supporting the connection between persistent pain sensitivity, reduced VTA 2-AG levels, and negative affect behaviors, with effects mitigated by exogenous 2-AG or retigabine treatment.

### pSNL Alters the VTA Expression of Kv7 Channels

eCB-related protein expression is influenced by eCB levels [56, 57]. Given reduced VTA 2-AG in pSNL mice and evidence that 2-AG may activate Kv7 channels [41], we analyzed Kv7 expression in VTA neurons using IHC. WK4 midbrain sections were stained with a Kv7.2 antibody, and a digital VTA overlay (Bregma: −3.16 mm to −3.88 mm) identified anatomical boundaries (**Fig. 2A**). Confocal analysis revealed significant cytosolic Kv7.2 expression within the VTA, characterized by staining in cell bodies and proximal basal dendrites (**Fig. 2B; Supplementary Figure 3**). DAPI staining clarified Kv7.2 cytosolic localization, consistent with previous reports [58].

**Fig. 2.**
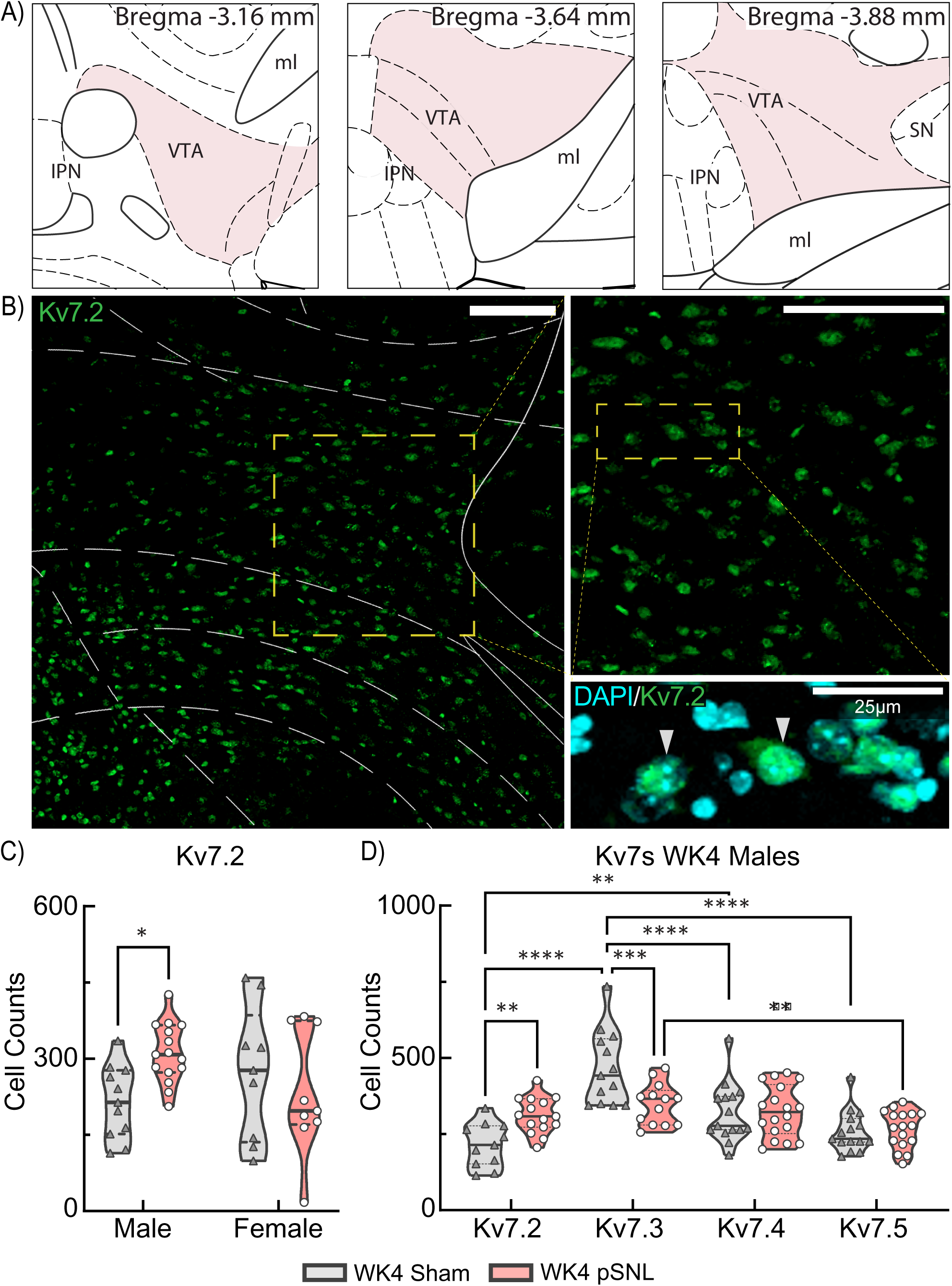
Sex-Specific Variations in VTA Kv7.2 Expression at WK4 pSNL. **a** Schematic representation of midbrain coronal brain sections adapted from “The Mouse Brain in Stereotaxic Coordinates” (Franklin & Paxinos, 2007, 3rd edition) illustrating evaluated VTA regions (*shaded*) at WK4 post-surgery. **b** Confocal photomicrographs of corresponding VTA sections from a WK4 sham control mouse, displaying immunostaining with Kv7.2 (*green*) in single-channel images at varying magnifications. Co-expression of Kv7.2+ with nuclear DAPI staining emphasizes cytosolic localization (see also **Supplementary Figure 3**). Images underwent rolling-ball background subtraction, linear contrast adjustment, and resizing for display; the DAPI channel was pseudocolored cyan. Scale bars: 100um (*left and right images*); 25 μm (*bottom image*). **c** Violin plots depict significant differences in Kv7.2 expression at WK4 based on sex and surgical group (interaction F(1,40)=5.441, *p=0.0248) revealing elevated Kv7.2 in male pSNL mice compared to sham controls (*p=0.0143). **d** A violin plot illustrates Kv7.2-5 subunit expression at WK4 following sham or pSNL treatment, with a 2W ME ANOVA indicating an interaction (F(3,78)=8.847, ****p<0.0001). pSNL was associated with increased Kv7.2 (**p=0.0026) and decreased Kv7.3 (***p=0.0006). Kv7.2 expression was significantly lower in sham controls compared to Kv7.3 (****p<0.0001) and Kv7.4 (**p=0.0034) at WK4 post-surgery, with no significant changes observed for Kv7.4 or Kv7.5 expression.

Further analysis of Kv7.2+ expression in WK4 mice demonstrated a significant sex-by-surgery interaction (*p=0.0248) (**Fig. 2C**). Fisher’s LSD post hoc analysis indicated a ∼1.5-fold increase in Kv7.2 expression in males (*p=0.0143), while females showed no significant change (p=0.3773), consistent with a sex-specific disparity after pSNL. Since Kv7 channels typically express as heteromeric combinations [59, 60], we utilized Kv7 subunit-specific antibodies for Kv7.3-7.5 and performed IHC in WK4 sham mice to evaluate overall Kv7 expression across VTA cells (**Fig. 2D**). As females showed no Kv7.2 expression changes, subsequent analysis of Kv7.3-7.5 was exclusive to male mice. As with Kv7.2, Kv7.3-5 displayed cytoplasmic staining patterns (**Supplementary Figure 3**). A 2W ME ANOVA confirmed a significant main effect of the subunit and subunit-by-surgery interaction (****p<0.0001) (**Fig. 2D**), indicating pSNL effects were specific to Kv7 subunits. Tukey’s post hoc analysis confirmed increased Kv7.2 (**p=0.0026) and decreased Kv7.3 (***p=0.0006) expression in pSNL mice, with no significant changes in Kv7.4 or Kv7.5 expression. These WK4 changes demonstrate critical pSNL effects in male mice on Kv7.2 and Kv7.3 subunits, primary components of inhibitory M-currents.

### Subunit Selective Changes in Kv7 Channels in VTA Dopamine Neurons

The VTA comprises multiple neuron types [61], raising the question of whether pSNL affected Kv7 subunit expression specifically within dopamine neurons. We employed double-label fluorescence IHC using tyrosine hydroxylase (TH) as a dopamine cell marker on WK4 tissue to assess this. Confocal analysis of co-labeled TH+Kv7.2-7.5 immunoreactive cells revealed strong cytosolic TH fluorescence within VTA neuronal somata in both sham and pSNL mice (**Fig 3A,B; Supplementary Figure 3**). Photomicrographs confirmed overall high co-expression of TH and Kv7.2-7.5, indicating subpopulations of VTA neurons with varying Kv7 expression (**Fig 3B; Supplementary Figure 3**). Quantitative analysis of TH expression showed similar numbers of TH+ neurons across both sexes and surgical groups (2W ME ANOVA, no significant differences). Based on observed sex-dependent Kv7.2 changes, we analyzed Kv7.2 co-expression with TH. A 2W ME ANOVA revealed that increased TH+Kv7.2 expression in pSNL mice resulted from a sex-dependent interaction (**p=0.0024). Fisher’s LSD post hoc analysis indicated significant increases in TH+Kv7.2 co-expression exclusively in male pSNL mice (****p=0.0001), while females showed no corresponding increase (p=0.2150), underscoring sex-specific differences in the VTA.

**Fig. 3.**
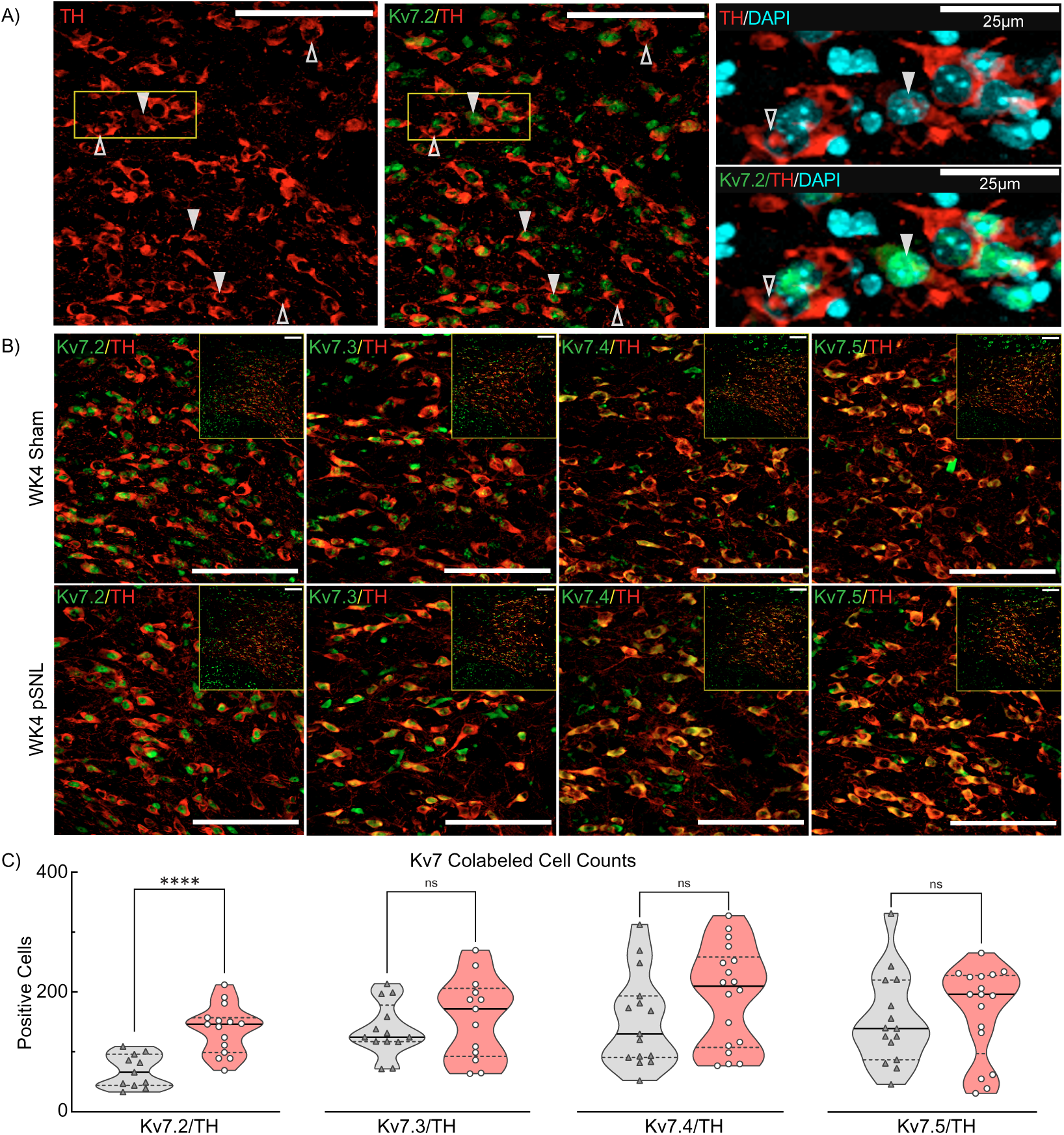
Elevated Kv7.2 Expression in TH+ VTA Dopamine Cells at WK4 pSNL. **a** Confocal photomicrographs of the contralateral VTA region from a male mouse at four weeks control sham-surgery, showing varying magnifications. Immunostaining reveals TH (*red*), Kv7.2 (*green*), and DAPI staining presented in single-channel images and overlays. Co-expression (TH+/Kv7.2+,▾) differentiates between cells expressing both markers versus those expressing only one. Scale bars: 100 μm; 25um. **b** Similar photomicrographs from WK4 male mice divided into sham-operated (*top panels*) and pSNL (*bottom panels*) groups, depicting co-localization of TH (*red*) with Kv7.2-5 (*green*). Images underwent rolling-ball background subtraction, linear contrast adjustment, and resizing for display. Scale bars: 100 µm. **c** Violin plots quantifying co-labeled VTA TH+ and Kv7.2-5+ cells. Increased Kv7.2 expression in pSNL mice compared to sham-operated controls was significant (t(24)=4.868. ****p<0.0001). No significant differences were detected for Kv7.3 (t(24)=0.8747 p=0.3904), Kv7.4 (t(31)=1.426 p=0.1638), or Kv7.5 expression (t(30)=0.5523, p=0.5849).

Quantification of co-expressed markers showed a two-fold increase in TH+Kv7.2 co-expression in pSNL mice (p<0.0001) (**Fig. 3C**). No significant changes were observed for TH with Kv7.3 (p=0.3904), Kv7.4 (p=0.1638), or Kv7.5 expression (p=0.5849) (**Fig. 3C**). DAPI staining showed no difference in overall VTA cell number (p=0.3397; not shown). Conversely, in sham-operated males, a 1W ANOVA identified significant differences in TH+Kv7 expression based on subunit (**p=0.0042). Specifically, TH+Kv7.2 expression was significantly lower than TH+Kv7.3 (−51.1%*, p=0.0378), TH+Kv7.4 (−55.9%, **p=0.0059), and TH+Kv7.5 (−55.1%, **p=0.0082), suggesting VTA dopamine neurons exhibit low basal Kv7.2 expression and a heightened expression during chronic neuropathic pain. Collectively, these multiple findings indicate that chronic pain and heightened negative affect-like behaviors in male mice are associated with reduced 2-AG and increased Kv7.2 expression specifically in VTA dopamine neurons. Decreased Kv7.3 expression likely occurs in non-dopamine (TH-) neurons, indicating cell type-specific Kv7 subunit adaptations in CPNA.

### Time-Dependent Changes in VTA Proteomic Landscape

Both 2-AG and Kv7 are crucial participants in common pathways underpinning neuronal development and maturation [62, 63]. To determine if pSNL pain alters interrelated pathway protein abundance in the VTA and to characterize these alterations over time, we conducted an unbiased proteomics screen comparing protein levels in pSNL and sham-operated mice at WK1 and WK4 (**Fig. 4A**). Pooled bilateral VTA punches from pairs of mice ensured adequate protein yield (WK1 sham, n=3; WK1 pSNL, n=4; WK4 sham, n=5; WK4 pSNL, n=5). Mass spectrometry identified 7 116 distinct proteins within the VTA proteome.

**Fig. 4.**
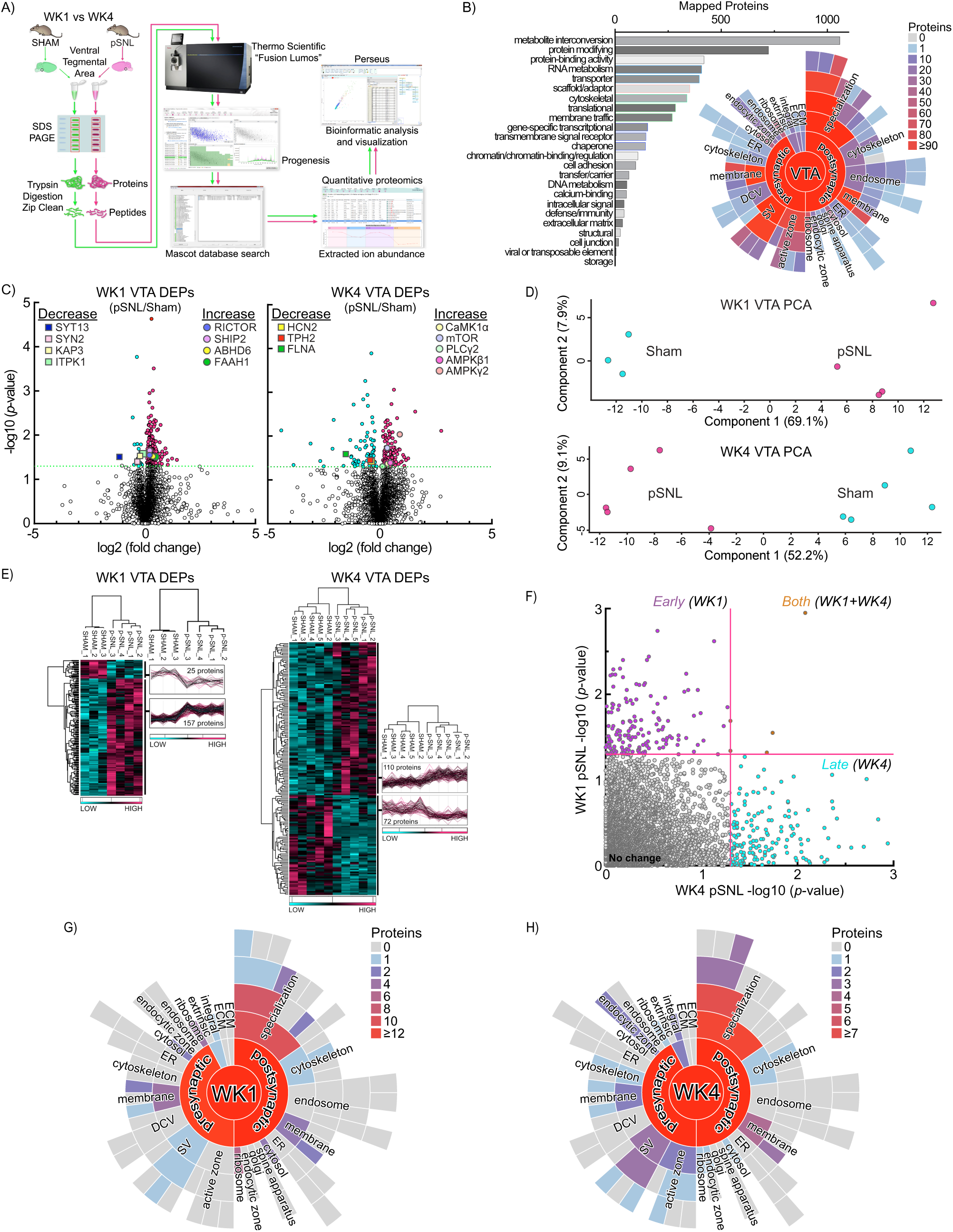
Comprehensive VTA Proteomic Analysis Following pSNL. **a** Workflow for the proteomic analysis comparison of VTA protein expression at WK1 and WK4 post-surgery. **b** PANTHER analysis categorized 5 251 proteins in the VTA proteome based on functional class. The Synaptic Gene Ontology (SynGO) starburst plot identified 1 203 proteins with defined subcellular localizations. Plots display the synapse (center), with pre- and postsynaptic sites in the first ring, and subsequent child terms in outer rings. **c** Volcano plots showing differentially expressed protein abundance (DEP) at WK1 and WK4, with significant differences indicated by −log10(*p*−value) ≥ 1.30. **d** Principal component analysis (PCA) and hierarchical clustering (heatmap) further validated significant separation between surgical cohorts at WK1 and WK4. **e** Heat maps from unbiased hierarchical clustering of 372 significantly affected proteins show reproducible abundance shifts. **f** Scatter plots demonstrate VTA proteome changes due to surgical intervention and time. Statistical differences are shown as −log10(*p*-value) for WK1 (*y-axis*) versus WK4 (*x-axis*). Red lines mark the p-value cutoff of < 0.05. Few proteins were changed at both timepoints, highlighting a differential VTA proteome at WK4, associated with CPNA, compared to WK1 pain. **g, h** The pSNL-enriched VTA proteome at WK1 (*g*) and WK4 (*h*) mapped to synaptic localizations using SynGO. Out of 372 identified DEPs in (*c*), 70 mapped to synaptic locations, including cytoskeleton and postsynaptic density (PSD). Protein quantities within each term are represented by a color scheme. Gray panels indicate SynGO terms without associated proteins; colored terms reflect changes in protein abundance in at least one module. Despite differing proteins affected by pSNL at WK1 and WK4, enrichment disparities predominantly occurred within overlapping cellular compartments.

PANTHER analysis categorized 5 251 VTA proteome proteins into functional classes, predominantly metabolite interconversion (1 059), protein-modifying (723), and protein-binding activity (419) (**Fig 4B**, *bar graph*). SynGO, a curated knowledgebase of synapse-specific proteins, identified 1 203 proteins corresponding to cellular components (CC), highlighting diverse synaptic localizations within the VTA (**Fig 4B**, starburst plot).

To track proteomic changes specific to timepoints, we compared protein abundance for pSNL and sham mice at WK1 and WK4 (**Fig 4C**, *left/right*), when negative affect-like behavior was apparent (**Fig. 1E**). We discerned 185 differentially expressed proteins (DEPs) at WK1 and 187 DEPs at WK4 (**Fig. 4C**; **Supplementary Table 2**). Both instances showed notable increases in protein abundance associated with pSNL treatments. Key WK4 DEPs included TPH2, PLCγ2, HCN2, mTOR, CaMK1α, AMPKγ2, and AMPKβ1. PCA (**Fig. 4D**) and hierarchical clustering (**Fig. 4E**) further validated significant separation between surgery groups with minimal within-group variance (7.9% at WK1, 9.1% at WK4) (**Fig 4D**). pSNL samples exhibited consistent differences in effective protein abundance relative to sham mice (**Fig 4E**).

Further comparison of DEPs between timepoints showed minimal overlap, indicating dynamic protein abundance shifts as chronic pain evolved (**Fig. 4F**). SynGO analysis explored synapse-specific subcellular localization for WK1 and WK4 (**Fig. 4G,H**). At WK1, 40 DEPs mapped to unique SynGO CC annotations and 27 to Biological Processes (BP) (not shown). At WK4, 30 DEPs mapped to CC annotations and 22 to BP. Both timepoints demonstrated protein abundance variations in presynaptic vesicles, membranes, ribosomes, postsynaptic densities (PSDs), and cytoskeleton proteins. These observations reveal that chronic pain-related proteomic alterations occur within similar synaptic domains but engage distinct proteins over time, aligning with established synaptic remodeling patterns [64, 65]. A substantial subset of WK4-detected proteins (CaMK1α, AMPKβ1, AMPKγ2, etc.) are associated with the CaMKK2 signaling pathway [20, 23, 66], implicated in intracellular calcium signaling alterations during synaptic remodeling [20, 25].

### Enrichment in Proteins Related to Dendritic Reorganization and CaMK1**α**

To investigate broader WK4 pSNL implications on VTA signaling, we performed gene ontology (GO) CC, BP, molecular function (MF), and Reactome pathway enrichment analyses of DEPs using DAVID (**Fig. 5A,B,C,D**). Analysis revealed significant enrichment in various GO CCs, MFs, BPs, and Reactome pathways (**Fig. 5A,B,C,D**). Notably, many enriched processes and functions relate to dendritic reorganization, including protein kinase binding, calmodulin binding, positive regulation of neuron projection development, and protein localization to the cell surface.

**Fig. 5.**
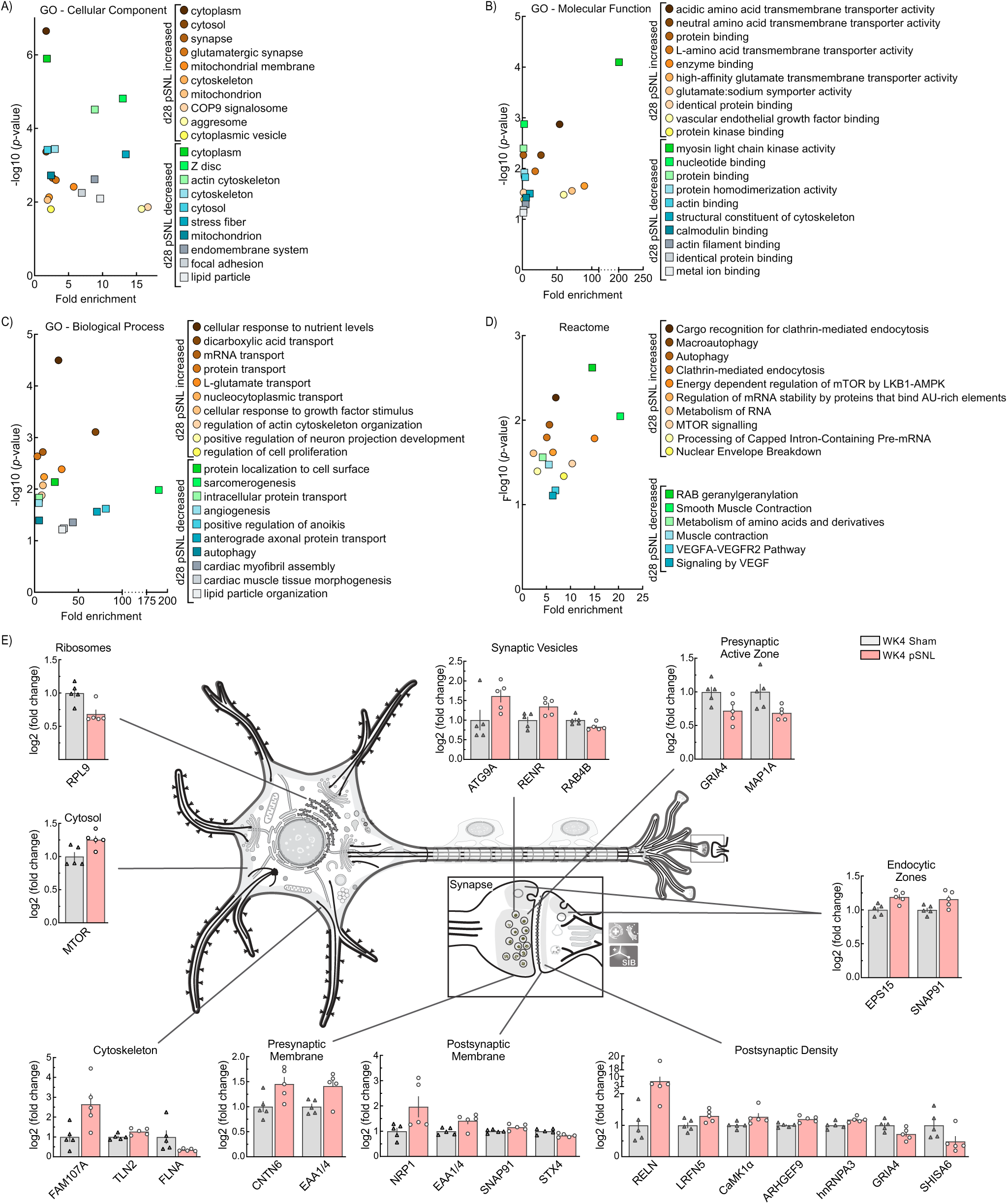
Changes in the Abundance of WK4 VTA Proteins. **a,b,c,d** VTA proteomic enrichment analysis using the DAVID tool identified significant gene ontology (GO) and reactome enrichment at WK4. Cellular components (CC), biological processes (BP), molecular functions (MF), and reactome pathways enriched from these proteins include roles in synaptogenesis, neuronal signaling, and transporter trafficking. **e** A neuronal illustration adapted from SwissBioPics depicts the subcellular localization and log2 fold changes of significantly altered synapse-specific proteins in WK4 pSNL mice. Bar graphs are expressed as mean ± SEM. Neuron illustration adapted from SwissBioPics: A resource for biological illustrations (https://www.swissbiopics.org/name/Animalneuronalcell).

In the comparative proteomic analysis, 15 synapse-specific (SynGO) WK4 DEPs showed increased abundance, while 7 exhibited decreased abundance (**Fig. 5E**). SynGO WK4 DEPs were localized to the cytosol, endocytic zones, and presynaptic membranes; decreases were primarily in presynaptic active zones and ribosomal structures. PSD, cytoskeleton, synaptic vesicles, and postsynaptic membranes displayed both increases and decreases. These results align with prior studies on dendritic reorganization [67].

Given CaMKK2’s potential involvement in dendritic reorganization, we specifically examined CaMK1α, a downstream kinase enhancing dendritic complexity via activity-dependent mechanisms [27–29]. Proteomic comparisons indicated significantly increased CaMK1α abundance in WK4 pSNL mice (**Fig 6A**, *p=0.0488). To validate, we employed IHC co-labeling to assess VTA CaMK1α expression. Utilizing TH (**Fig. 6B**, *red*) and CaMK1α (**Fig. 6B**, *green*) on VTA slices from WK4 sham and pSNL mice, double-labeling revealed distinct somatodendritic staining patterns with sporadic co-localization. Statistical analyses confirmed significant increases in CaMK1α expression (*p=0.0253) and TH+CaMK1α co-labeling (**p=0.0018) in WK4 pSNL mice (**Fig. 6C**).

**Fig 6.**
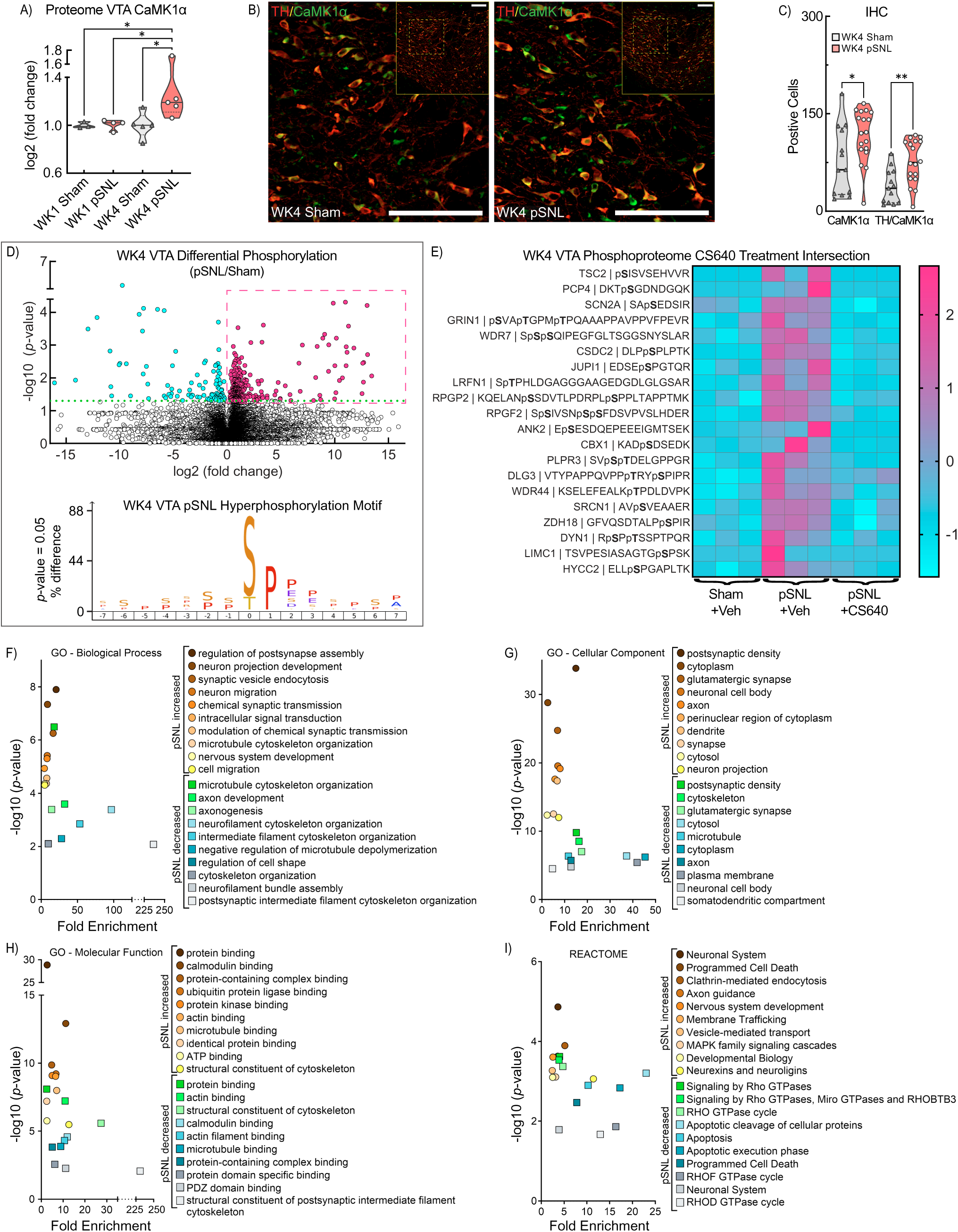
Activation of VTA CaMKK2-CaMK1α Axes. ***a*** Violin plot illustrating CaMK1α abundance in the VTA proteome, normalized to WK1 sham mice. WK4 pSNL mice show elevated CaMK1α (1W ANOVA: F(3,13) = 3.441, *p* = 0.0488; WK1 Sham vs. WK4 pSNL, *p* = 0.0348; WK1 pSNL vs. WK4 pSNL, *p* = 0.0257; WK4 Sham vs. WK4 pSNL, *p* = 0.0175). ***b*** Cropped confocal photomicrograph shows TH+ (*red*) and CaMK1α+ (*green*) in VTA neurons. Inset shows full VTA photomicrograph for the same male WK4 sham (*left*) and pSNL (*right*) mice. Images underwent rolling-ball background subtraction, linear contrast adjustment, and resizing for display. ***c*** Violin plot displays VTA cell counts for CaMK1α+ and TH+CaMK1α in WK4 sham and pSNL mice (unpaired t-tests: t(29) = 2.359, \**p* = 0.0253; t(29) = 3.434, \*\**p* = 0.0018; solid lines in violin plots denote the median, while dashed lines indicate quartiles). ***d*** *(top)* Volcano plot illustrating the effect of pSNL on VTA-specific phosphorylated peptides at WK4 (n=3/samples per group), with significant differences expressed as −log10(p-value) ≥ 1.30. The y-axis reflects −log10 adjusted *p* values; blue symbols represent decreased abundance of phosphopeptide ions while red symbols and box indicate increased abundance in pSNL compared to sham. ***d*** *(bottom)* iceLogo of hyperphosphorylation motif in WK4 pSNL mice illustrate amino acids at phosphosites with p-values < 0.05. ***e*** Heat map shows unbiased hierarchical clustering of 20 significantly increased phosphorylated peptides at WK4 in pSNL versus sham-operated mice, is reduced following CS640 treatment (2 μg/μL, ICV) in pSNL versus CS640-treated pSNL mice, confirming consistent expression patterns across biological samples (9 mice; 3 per sample). ***f****, **g, h, i*** VTA phosphoproteomic enrichment analysis utilizing the DAVID identifying significant increases and decreases at WK4 post-pSNL. Key biological processes (BP) (*f)*, cellular components (CC) (*g*), molecular functions (MF) (*h)*, and reactome pathways (*i)* significantly enriched including multiple proteins associated with the CaMKK2-CaMK1α axis linked to dendritic morphogenesis.

### pSNL Changes in the VTA Phosphoproteome

To further evaluate chronic pain’s effect on CaMK1α activity levels, we employed phosphoproteomic analysis. Pooled bilateral tissue punches from three mice per group (WK4 vehicle-treated sham and pSNL) yielded 29 605 phosphopeptide ions associated with 2 676 unique proteins (**Fig. 6D**, *top,* volcano plot). Comparing phosphorylation states between vehicle-treated pSNL and sham groups, we identified significant alterations in 577 phosphopeptide ions, with a predominance of hyperphosphorylation in pSNL mice relative to sham controls (**Fig. 6D**, *top, red+blue circles***; Supplementary Table 2**). These alterations included differential phosphorylation of proteins integral to the CaMKK2 signaling pathway and Kv7.2 modulation, such as CaMKK2, HCN2, ANK2, PCP4, RYR2, and CaMKIIα (**Supplementary Table 2**) [20, 23, 66, 68–71]. Motif analysis of the hyperphosphorylated sites in pSNL (Fig 6D, *top, red box*) with iceLogo revealed proline enriched targets (**Fig. 6D**, *bottom*). To corroborate these findings with our CaMK1α discovery, we used a next-generation CaMK1 blocker (CS640) and quantified the VTA phosphoproteome of CS640-treated pSNL mice (2 μg/μl, ICV). Among phosphopeptides upregulated in pSNL and downregulated with CS640, we identified 20 proteins across 28 phosphorylation sites (**Fig. 6E**). GO and Reactome pathway enrichment analyses of the proteins with significantly affected phosphorylation in vehicle-treated pSNL mice noted relevant biological categories, including axonogenesis, regulation of postsynapse assembly, PSDs, calmodulin binding, protein kinase binding, and clathrin-mediated endocytosis (**Fig. 6F,G,H,I**). These studies indicated the potential for an increased role for CaMK1α, which, alongside other VTA kinases, may contribute to the CPNA phenotype [25]. Collectively, these data link sustained pSNL pain and associated negative affect-like behavior with temporal alterations in 2-AG, Kv7.2 channels, and the VTA proteomic landscape. These changes, characterized by enhanced activity in the VTA CaMKK2-CaMK1α/−AMPK axis, support the notion of 2-AG, Kv7, and dendritic signaling reorganization during CPNA.

## Discussion

Chronic Pain Negative Affect (CPNA) is a debilitating comorbidity of chronic pain that complicates patient management and necessitates new therapeutic strategies. Our study advances translational efforts by providing fundamental neurobiological insights and identifying actionable molecular targets with preclinical therapeutic potential. We investigated CPNA mechanisms in the VTA—a critical brain region for reward, mood, and pain processing.

By employing a multidisciplinary approach that combined immunohistochemistry with lipidomic, proteomic, and phosphoproteomic screening in a pSNL mouse model, we generated an unprecedented molecular map of the VTA during CPNA progression. While previous studies have used targeted analyses or omic screening in the VTA [9, 72, 73] and profiled broader brain regions [74–76], our integrated multi-omic strategy offers a uniquely deep and unbiased characterization. We identified specific molecular alterations within VTA dopamine neurons, including sex-dependent Kv7.2 channel upregulation in males and profound changes in 2-AG levels and the CaMKK2-CaMK1α/−AMPK pathways. We then demonstrated the therapeutic actionability of these nodes through preclinical interventions. We hypothesize that these concurrent dysregulations drive a pathological cascade leading to VTA dopamine neuron hyperexcitability and the persistent negative affective states of CPNA, a concept consistent with existing literature [7–11]. Our translational findings show that both exogenous 2-AG and the Kv7 channel opener retigabine effectively alleviated pain-related negative affect, representing a conceptual advance in understanding CPNA pathophysiology.

While robust, our findings in an animal model require future clinical validation. Our investigation at one and four weeks post-injury offers crucial insights into early adaptations, but broader temporal studies may reveal additional dynamics. Furthermore, CPNA involves interconnected neural circuitry beyond the VTA [74–77]. A key unresolved issue is the conflicting reports on VTA dopamine neuron activity in chronic pain. While our findings implicate mechanisms that may underlie the hyperexcitability reported in some studies [7–11], other research indicates that inflammatory pain can decrease VTA dopamine neuron activity, leading to anhedonia [11–18]. These discrepancies may stem from differences in timing, pain models, VTA subpopulations, or behavioral assays, which our study did not resolve. Nevertheless, our results provide several key observations.

### Endocannabinoid System (ECS) and 2-AG Dysregulation

One prominent system affected in CPNA is the ECS, particularly the lipid mediator 2-AG. We found significant reductions in VTA 2-AG levels in male pSNL mice, which paralleled negative affect-like behaviors. This aligns with human studies showing reduced plasma 2-AG in negative affect [35–37] and preclinical work demonstrating disrupted 2-AG metabolism in anxiety and other negative affective states [74, 78]. Crucially, our finding that exogenous 2-AG attenuates CPNA symptoms—similarly to retigabine [9, 10]—directly links 2-AG deficiency to symptomology and underscores its therapeutic potential. This is supported by literature showing that targeting 2-AG metabolism via MAGL inhibition has antidepressant/anxiolytic effects in mice [79] and reduces pain in rats [53]. Further evidence includes 2-AG’s protective effects against LPS-induced impairments in vitro [80] and the observation that rTMS therapy increases hippocampal 2-AG in rats [81].

Beyond classical receptor agonism, eCBs can directly modulate ion channels, with 2-AG acting as a positive allosteric modulator of Kv7.2 [41]. Evidence also suggests 2-AG can indirectly alter AMPK via CB1R-CaMKK-AMPK signaling in cardiomyocytes [39]. Other less-reported, CB1-independent actions of 2-AG involve platelets [82] and other non-CB1 proteins and ion channels (i.e., TRPV, N-type VGCC) [83]. Similarly, anandamide (AEA), which was not impacted in our study, also exerts non-canonical actions at targets like TRPV, GPR55, GPR18, and PPARs [84, 85].

While enhancing VTA 2-AG shows therapeutic promise for CPNA, the role of 2-AG in pain is complex. For instance, excessive circulating 2-AG at the time of injury may paradoxically increase chronic pain risk [32]. Furthermore, pain is often comorbid with cognitive impairments involving the ECS [86], and hormonal cycles can influence ECS function and cannabinoid responsiveness [87]. These points highlight the need to distinguish between acute injury responses and chronic pain states, as well as systemic versus brain-region-specific effects and the significant role of sex.

### Kv7.2 Channel Dysregulation and Sex-Dependent Effects

The dysregulation of Kv7.2 channels adds another critical layer to VTA dysfunction in CPNA. While neuronal hyperexcitability in pain is associated with decreased Kv7 channel expression in the DRG [88], our data reveal a paradoxical, sex-specific upregulation of Kv7.2 protein, but not Kv7.3, in VTA dopamine neurons of male pSNL mice. This counterintuitive increase suggests that functional modification, rather than expression level, may be the primary driver of dysfunction, contributing to hyperexcitability despite greater protein abundance. The critical role of Kv7.2 in regulating neuronal excitability is underscored by evidence that its removal alone can trigger spreading depression [89]. Moreover, pathogenic loss-of-function variants cause severe neurodevelopmental disorders [62], highlighting the channel’s fundamental importance.

Increased Kv7.2:Kv7.3 ratios can produce less functional homomeric Kv7.2 channels with weaker M-currents [90, 91]. Our observed sex-dependent upregulation points to distinct mechanisms underlying CPNA symptomology in males and females. The C-terminal tails of Kv7 channels, which are critical for trafficking, are targets for kinases, calmodulin, and PIP2 [92, 93]. Multiple kinases phosphorylate Kv7 channels, including CaMKII, PKA, PKC, and MAP kinases [94]. Our finding of increased CaMKK2 phosphorylation supports such kinase-mediated Kv7.2 dysfunction via several plausible pathways: CaMKK activates MAP kinases that regulate Kv7 [94, 95]; PKA directly modulates Kv7.2 [96]; and crosstalk between CaM kinases and PKC decreases Kv7.2 channel activity [95].

### CaMKK2-CaMK1α/−AMPK Pathways

Our multi-omic analyses identify the CaMKK2 signaling cascade as a central node in the VTA’s adaptation to CPNA. We observed increased CaMKK2 phosphorylation, which we hypothesize is a maladaptive homeostatic response to pain-induced VTA hyperexcitability, consistent with principles of cellular plasticity under chronic stress [97–99]. Despite inhibitory phosphorylation of CaMKK2 at S495, the absence of S100/S511 phosphosites suggests residual activity, which may sustain some downstream signaling while hyperexcitable neurons compensate by increasing downstream kinase subunits like AMPK and CaMK1α. The increased abundance of AMPKγ2 in CPNA suggests an attempt to compensate for metabolic stress, which could amplify VTA dysregulation. AMPK influences both protein and lipid synthesis and degradation, as well as ion channel trafficking [22, 23, 25, 26, 66].

Similarly, dysregulation of CaMK1α, critical for structural plasticity and spine density [30], may impair the VTA’s ability to adapt to pain stressors, potentially leading to excessive immature synapses and perpetuating hyperexcitability [100]. The interplay between CaMK1α and AMPK, collectively influencing dendritic reorganization and synaptic integrity, appears to be a key mechanism shaping the VTA’s maladaptive response. While direct evidence linking CaMKK2 to endocannabinoid-metabolizing enzymes like MAGL or DAGL is lacking, its regulation of cellular energy and signaling cascades intertwined with lipid metabolism is well-established [20, 22, 23, 38]. It is important here to also note we found that the CaMK1α antagonist CS640 reversed key pSNL-related changes in phosphorylated proteins involved in morphological rearrangement, providing compelling pharmacological evidence for this pathway as an actionable therapeutic target.

## Conclusion

This study provides the first comprehensive, multi-omic analyses of molecular adaptations in the VTA linked to CPNA. By identifying critical neurobiological substrates—including dysregulation in Kv7.2 channels, the endocannabinoid system, and the CaMKK2 pathway—and elucidating their interplay, we provide a robust foundation for developing targeted therapies. The sex-dependent nature of our findings underscores the importance of sex as a biological variable in CPNA research.

Future work should focus on how Kv7.2 dysregulation and 2-AG reduction mechanistically contribute to VTA hyperexcitability and negative affect. Investigating the functional consequences of CaMKK2 pathway alterations and their interplay with Kv7.2 and 2-AG is paramount. Long-term translational studies are warranted to assess the sustained therapeutic potential of Kv7.2 modulators, 2-AG enhancers, and CaMK1α antagonists. Finally, rigorous delineation of the specific cannabinoid receptor mechanisms and further investigation into sex-specific signaling differences are crucial next steps to advance innovative and personalized strategies aimed at alleviating the burden of CPNA.

## Acknowledgements

This work was funded by grants from the National Institute on Drug Abuse through the University of Arizona Center of Excellence in Addiction Studies (CEAS; P30DA051355), with additional support from the Department of Pharmacology Comprehensive Center for Pain and Addiction (CCPA) in the College of Medicine at the University of Arizona. The University of Arizona Cancer Center Analytical Chemistry Shared Resources (ACSR) is acknowledged for assistance with LC-MS services. The University of Arizona Health Sciences Quantitative Proteomics Laboratory is acknowledged for MS-based proteomic and phosphoproteomic support.

## Author Contributions

C.C.D., L.A.M., P.R.L., T.W.V., A.C.R. contributed to the design of the study. All authors participated in the acquisition of data. C.C.D., L.A.M., S.M.D.B., P.R.L., A.C.R. contributed to the analysis of the data. C.C.D., P.R.L., A.C.R. contributed to the interpretation of the data. C.C.D. and A.C.R. wrote the initial draft of the paper. All authors revised the paper for intellectual content and provided final approval of the paper to be published.

---

**Supplementary Figure 1. pSNL surgery causes long-lasting allodynia.** Violin plot depicts individual animal paw withdrawal threshold values in male and female mice from sham and pSNL-exposed groups (interaction F_(2,151)_=39.46, *\*\*\*\*p*<0.0001). Both groups exhibited hypersensitivity at WK1 compared to baseline (*\*\*\*\*p*<0.0001) to different extents (*\*\*p*=0.0030). While pSNL mice continued to show hypersensitivity at WK4 (*\*\*p*<0001), sham-exposed mice returned to withdrawal thresholds similar to baseline *\*\*\*\*p*<0.0001). Sample sizes are indicated in parentheses (Sham male, Sham female; pSNL male, pSNL female). Solid lines in violin plots depict median and dashed lines depict quartiles.

**Supplementary Figure 2. Early Imbalance in VTA Neuromodulation at WK1 pSNL.** Bar graph depicts the relative abundance of various proteins normalized to sham-operated mice (multiple unpaired t-tests). Included are enzymes involved in endocannabinoid (eCB) metabolism (fatty acid amide hydrolase (FAAH) and alpha/beta-hydrolase domain containing 6 (ABHD6)) and eCB synthesis (monoacylglycerol lipase (MGLL) and diacylglycerol lipase alpha (DAGLA)). Also analyzed are proteins important for anandamide production (N-acyl-phosphatidylethanolamine phospholipase D (NAPEPLD), and experience dependent synaptic plasticity (KS6KA5 (MSK1), and KAP3 (Prkar2b)). All data are expressed as mean ± SEM.

**Supplementary Figure 3. Cytosolic Expression Patterns of TH+ and Kv7.2-5 Among VTA cells.** High magnification confocal photomicrographs of the contralateral VTA region from male mice, divided into sham-operated (*left two panels*) and pSNL (*right panel*) groups, at four weeks post-surgery. Images depict co-localization of TH (*red*), Kv7.2-5 (*green*, presented in descending order), and nuclear DAPI staining (*blue*). Magnified images focusing on individual neurons (*far right*) show only Kv7 and DAPI staining. Images underwent rolling-ball background subtraction, linear contrast adjustment, and resizing for display. Scale bars represent 25 µm and asterisks represent the nucleus.

## References

1. Treede R-D, Rief W, Barke A, Aziz Q, Bennett MI, Benoliel R, et al. A classification of chronic pain for ICD-11. PAIN. 2015;156:1003.

2. De La Rosa JS, Brady BR, Ibrahim MM, Herder KE, Wallace JS, Padilla AR, et al. Co-occurrence of chronic pain and anxiety/depression symptoms in U.S. adults: prevalence, functional impacts, and opportunities. PAIN. 2024;165:666.

3. Hooten WM. Chronic Pain and Mental Health Disorders: Shared Neural Mechanisms, Epidemiology, and Treatment. Mayo Clinic Proceedings. 2016;91:955–970.

4. Probst T, Neumeier S, Altmeppen J, Angerer M, Loew T, Pieh C. Depressed Mood Differentially Mediates the Relationship between Pain Intensity and Pain Disability Depending on Pain Duration: A Moderated Mediation Analysis in Chronic Pain Patients. Pain Res Manag. 2016;2016:3204914.

5. Uebelacker LA, Weisberg RB, Herman DS, Bailey GL, Pinkston-Camp MM, Stein MD. Chronic Pain in HIV-Infected Patients: Relationship to Depression, Substance Use, and Mental Health and Pain Treatment. Pain Med. 2015;16:1870–1881.

6. Hou G, Hao M, Duan J, Han M-H. The Formation and Function of the VTA Dopamine System. IJMS. 2024;25:3875.

7. Morris LS, Costi S, Hameed S, Collins KA, Stern ER, Chowdhury A, et al. Effects of KCNQ potassium channel modulation on ventral tegmental area activity and connectivity in individuals with depression and anhedonia. Mol Psychiatry. 2025. 25 March 2025. 10.1038/s41380-025-02957-7.

8. Friedman AK, Juarez B, Ku SM, Zhang H, Calizo RC, Walsh JJ, et al. KCNQ channel openers reverse depressive symptoms via an active resilience mechanism. Nat Commun. 2016;7:11671.

9. Zhang L, Ji M, Sun Y, Wang Q, Jin M, Wang S, et al. VTA dopaminergic neurons involved in chronic spared nerve injury pain-induced depressive-like behavior. Brain Research Bulletin. 2025;222:111261.

10. Zhang L, Wang J, Niu C, Zhang Y, Zhu T, Huang D, et al. Activation of parabrachial nucleus - ventral tegmental area pathway underlies the comorbid depression in chronic neuropathic pain in mice. Cell Reports. 2021;37:109936.

11. Fox ME, Lobo MK. The molecular and cellular mechanisms of depression: a focus on reward circuitry. Mol Psychiatry. 2019;24:1798–1815.

12. Wang X-Y, Jia W-B, Xu X, Chen R, Wang L-B, Su X-J, et al. A glutamatergic DRN–VTA pathway modulates neuropathic pain and comorbid anhedonia-like behavior in mice. Nat Commun. 2023;14:5124.

13. Flores-García M, Rizzo A, Garçon-Poca MZ, Fernández-Dueñas V, Bonaventura J. Converging circuits between pain and depression: the ventral tegmental area as a therapeutic hub. Front Pharmacol. 2023;14.

14. Barker DJ, Zhang S, Wang H, Estrin DJ, Miranda-Barrientos J, Liu B, et al. Lateral preoptic area glutamate neurons relay nociceptive information to the ventral tegmental area. Cell Rep. 2023;42:113029.

15. Ji Y, Xiao Y, Bai X, Gu J, Ma T, Feng Y, et al. Modulation of comorbid depression of neuropathic pain by dopamine input from VTA to the ventral hippocampus. Theranostics. 2025;15:4101–4123.

16. Song Q, Wei A, Xu H, Gu Y, Jiang Y, Dong N, et al. An ACC-VTA-ACC positive-feedback loop mediates the persistence of neuropathic pain and emotional consequences. Nat Neurosci. 2024;27:272–285.

17. Huang S, Borgland SL, Zamponi GW. Peripheral nerve injury-induced alterations in VTA neuron firing properties. Molecular Brain. 2019;12:89.

18. Markovic T, Pedersen CE, Massaly N, Vachez YM, Ruyle B, Murphy CA, et al. Pain induces adaptations in ventral tegmental area dopamine neurons to drive anhedonia-like behavior. Nat Neurosci. 2021;24:1601–1613.

19. Wang J, Li Y. KCNQ potassium channels in sensory system and neural circuits. Acta Pharmacol Sin. 2016;37:25–33.

20. Tokumitsu H, Sakagami H. Molecular Mechanisms Underlying Ca2+/Calmodulin-Dependent Protein Kinase Kinase Signal Transduction. Int J Mol Sci. 2022;23:11025.

21. Elzière L, Sar C, Ventéo S, Bourane S, Puech S, Sonrier C, et al. CaMKK-CaMK1a, a New Post-Traumatic Signalling Pathway Induced in Mouse Somatosensory Neurons. PLoS ONE. 2014;9:e97736.

22. Wang S, Dai Y. Roles of AMPK and Its Downstream Signals in Pain Regulation. Life (Basel). 2021;11:836.

23. McAloon LM, Muller AG, Nay K, Lu EL, Smeuninx B, Means AR, et al. CaMKK2: bridging the gap between Ca2+ signaling and energy-sensing. Essays in Biochemistry. 2024;68:309–320.

24. Chen L, Ni C, Lu D, Zhang S, Li Y, Wang D, et al. Curcumin analog C16 attenuates bone cancer pain induced by MADB 106 breast cancer cells in female rats and inhibits the CREB/NLGN2 signaling axis by targeting CaMKⅠα. Neuropharmacology. 2025;266:110284.

25. Kaiser J, Nay K, Horne CR, McAloon LM, Fuller OK, Muller AG, et al. CaMKK2 as an emerging treatment target for bipolar disorder. Mol Psychiatry. 2023;28:4500–4511.

26. Lee A, Kondapalli C, Virga DM, Lewis TL, Koo SY, Ashok A, et al. Aβ42 oligomers trigger synaptic loss through CAMKK2-AMPK-dependent effectors coordinating mitochondrial fission and mitophagy. Nat Commun. 2022;13:4444.

27. Ghiretti AE, Kenny K, Marr MT, Paradis S. CaMKII-Dependent Phosphorylation of the GTPase Rem2 Is Required to Restrict Dendritic Complexity. J Neurosci. 2013;33:6504–6515.

28. Wayman GA, Impey S, Marks D, Saneyoshi T, Grant WF, Derkach V, et al. Activity-Dependent Dendritic Arborization Mediated by CaM-Kinase I Activation and Enhanced CREB-Dependent Transcription of Wnt-2. Neuron. 2006;50:897–909.

29. Ageta-Ishihara N, Takemoto-Kimura S, Nonaka M, Adachi-Morishima A, Suzuki K, Kamijo S, et al. Control of Cortical Axon Elongation by a GABA-Driven Ca2+/Calmodulin-Dependent Protein Kinase Cascade. J Neurosci. 2009;29:13720–13729.

30. Fortin DA, Davare MA, Srivastava T, Brady JD, Nygaard S, Derkach VA, et al. Long-term potentiation-dependent spine enlargement requires synaptic Ca2+-permeable AMPA receptors recruited by CaM-kinase I. J Neurosci. 2010;30:11565–11575.

31. Fromont C, Atzori A, Kaur D, Hashmi L, Greco G, Cabanillas A, et al. Discovery of Highly Selective Inhibitors of Calmodulin-Dependent Kinases That Restore Insulin Sensitivity in the Diet-Induced Obesity in Vivo Mouse Model. J Med Chem. 2020;63:6784–6801.

32. Trevino CM, Hillard CJ, Szabo A, deRoon-Cassini TA. Serum Concentrations of the Endocannabinoid, 2-Arachidonoylglycerol, in the Peri-Trauma Period Are Positively Associated with Chronic Pain Months Later. Biomedicines. 2022;10:1599.

33. Fitzgerald JM, Chesney SA, Lee TS, Brasel K, Larson CL, Hillard CJ, et al. Circulating endocannabinoids and prospective risk for depression in trauma-injury survivors. Neurobiology of Stress. 2021;14:100304.

34. Mecca CM, Chao D, Yu G, Feng Y, Segel I, Zhang Z, et al. Dynamic Change of Endocannabinoid Signaling in the Medial Prefrontal Cortex Controls the Development of Depression after Neuropathic Pain. J Neurosci. 2021:JN-RM-3135-20.

35. Bersani G, Pacitti F, Iannitelli A, Caroti E, Quartini A, Xenos D, et al. Inverse correlation between plasma 2-arachidonoylglycerol levels and subjective severity of depression. Human Psychopharmacology: Clinical and Experimental. 2021;36:e2779.

36. Silveira KM, Wegener G, Joca SRL. Targeting 2-arachidonoylglycerol signalling in the neurobiology and treatment of depression. Basic Clin Pharmacol Toxicol. 2021;129:3–14.

37. Hillard CJ, Weinlander KM, Stuhr KL. Contributions of endocannabinoid signaling to psychiatric disorders in humans: genetic and biochemical evidence. Neuroscience. 2012;204:207–229.

38. Signorello MG, Leoncini G. Activation of CaMKKβ/AMPKα pathway by 2-AG in human platelets. J Cell Biochem. 2018;119:876–884.

39. Chanda D, Oligschlaeger Y, Geraets I, Liu Y, Zhu X, Li J, et al. 2-Arachidonoylglycerol ameliorates inflammatory stress-induced insulin resistance in cardiomyocytes. J Biol Chem. 2017;292:7105–7114.

40. Forner-Piquer I, Giommi C, Sella F, Lombó M, Montik N, Dalla Valle L, et al. Endocannabinoid System and Metabolism: The Influences of Sex. Int J Mol Sci. 2024;25:11909.

41. Incontro S, Sammari M, Azzaz F, Inglebert Y, Ankri N, Russier M, et al. Endocannabinoids Tune Intrinsic Excitability in O-LM Interneurons by Direct Modulation of Postsynaptic Kv7 Channels. J Neurosci. 2021;41:9521–9538.

42. Zhang H-XB, Heckman L, Niday Z, Jo S, Fujita A, Shim J, et al. Cannabidiol activates neuronal Kv7 channels. eLife. 2022;11:e73246.

43. Hiniesto-Iñigo I, Castro-Gonzalez LM, Corradi V, Skarsfeldt MA, Yazdi S, Lundholm S, et al. Endocannabinoids enhance hKV7.1/KCNE1 channel function and shorten the cardiac action potential and QT interval. EBioMedicine. 2023;89:104459.

44. Korah HE, Cheng K, Washington SM, Flowers ME, Stratton HJ, Patwardhan A, et al. Partial Sciatic Nerve Ligation: A Mouse Model of Chronic Neuropathic Pain to Study the Antinociceptive Effect of Novel Therapies. J Vis Exp. 2022. 6 October 2022. 10.3791/64555.

45. Martin LF, Almuslim M, Ismail KA, Ibrahim MM, Moutal A, Cheng K, et al. The conotoxin Contulakin-G reverses hypersensitivity observed in rodent models of cancer-induced bone pain without inducing tolerance or motor disturbance. Pain. 2025;166:376–387.

46. Parrilla-Carrero J, Buchta WC, Goswamee P, Culver O, McKendrick G, Harlan B, et al. Restoration of Kv7 Channel-Mediated Inhibition Reduces Cued-Reinstatement of Cocaine Seeking. J Neurosci. 2018;38:4212–4229.

47. Kruse R, Krantz J, Barker N, Coletta RL, Rafikov R, Luo M, et al. Characterization of the CLASP2 Protein Interaction Network Identifies SOGA1 as a Microtubule-Associated Protein. Mol Cell Proteomics. 2017;16:1718–1735.

48. Parker SS, Krantz J, Kwak E-A, Barker NK, Deer CG, Lee NY, et al. Insulin Induces Microtubule Stabilization and Regulates the Microtubule Plus-end Tracking Protein Network in Adipocytes. Mol Cell Proteomics. 2019;18:1363–1381.

49. Vizcarra VS, Barber KR, Franca-Solomon G, Majuta L, Smith A, Langlais PR, et al. Targeting 5-HT2A receptors and Kv7 channels in PFC to attenuate chronic neuropathic pain in rats using a spared nerve injury model. Neurosci Lett. 2022;789:136864.

50. Keresztes A, Olson K, Nguyen P, Lopez-Pier MA, Hecksel R, Barker NK, et al. Antagonism of the mu-delta opioid receptor heterodimer enhances opioid antinociception by activating Src and calcium/calmodulin-dependent protein kinase II signaling. Pain. 2022;163:146–158.

51. Levine AA, Liktor-Busa E, Balasubramanian S, Palomino SM, Burtman AM, Couture SA, et al. Depletion of Endothelial-Derived 2-AG Reduces Blood-Endothelial Barrier Integrity via Alteration of VE-Cadherin and the Phospho-Proteome. Int J Mol Sci. 2023;25:531.

52. Murataeva N, Straiker A, Mackie K. Parsing the players: 2-arachidonoylglycerol synthesis and degradation in the CNS. British J Pharmacology. 2014;171:1379–1391.

53. Liktor-Busa E, Levine AA, Palomino SM, Singh S, Wahl J, Vanderah TW, et al. ABHD6 and MAGL control 2-AG levels in the PAG and allodynia in a CSD-induced periorbital model of headache. Front Pain Res (Lausanne). 2023;4:1171188.

54. Yuan X-C, Zhu B, Jing X-H, Xiong L-Z, Wu C-H, Gao F, et al. Electroacupuncture Potentiates Cannabinoid Receptor-Mediated Descending Inhibitory Control in a Mouse Model of Knee Osteoarthritis. Front Mol Neurosci. 2018;11:112.

55. Stensson N, Ghafouri N, Ernberg M, Mannerkorpi K, Kosek E, Gerdle B, et al. The Relationship of Endocannabinoidome Lipid Mediators With Pain and Psychological Stress in Women With Fibromyalgia: A Case-Control Study. J Pain. 2018;19:1318–1328.

56. Medina-Saldivar C, Pardo GVE, Pacheco-Otalora LF. Effect of MCH1, a fatty-acid amide hydrolase inhibitor, on the depressive-like behavior and gene expression of endocannabinoid and dopaminergic-signaling system in the mouse nucleus accumbens. Braz J Med Biol Res. 2024;57:e12857.

57. Baggelaar MP, Maccarrone M, van der Stelt M. 2-Arachidonoylglycerol: A signaling lipid with manifold actions in the brain. Progress in Lipid Research. 2018;71:1–17.

58. Verneuil J, Brocard C, Trouplin V, Villard L, Peyronnet-Roux J, Brocard F. The M-current works in tandem with the persistent sodium current to set the speed of locomotion. PLOS Biology. 2020;18:e3000738.

59. Li L, Sun H, Ding J, Niu C, Su M, Zhang L, et al. Selective targeting of M-type potassium Kv7.4 channels demonstrates their key role in the regulation of dopaminergic neuronal excitability and depression-like behaviour. British J Pharmacology. 2017;174:4277–4294.

60. Liu E, Pang K, Liu M, Tan X, Hang Z, Mu S, et al. Activation of Kv7 channels normalizes hyperactivity of the VTA-NAcLat circuit and attenuates methamphetamine-induced conditioned place preference and sensitization in mice. Mol Psychiatry. 2023;28:5183–5194.

61. Morales M, Margolis EB. Ventral tegmental area: cellular heterogeneity, connectivity and behaviour. Nat Rev Neurosci. 2017;18:73–85.

62. Dirkx N, Miceli F, Taglialatela M, Weckhuysen S. The Role of Kv7.2 in Neurodevelopment: Insights and Gaps in Our Understanding. Front Physiol. 2020;11.

63. Rodrigues RJ, Marques JM, Köfalvi A. Cannabis, Endocannabinoids and Brain Development: From Embryogenesis to Adolescence. Cells. 2024;13:1875.

64. Van Oostrum M, Campbell B, Seng C, Müller M, Tom Dieck S, Hammer J, et al. Surfaceome dynamics reveal proteostasis-independent reorganization of neuronal surface proteins during development and synaptic plasticity. Nat Commun. 2020;11:4990.

65. Heo S, Kang T, Bygrave AM, Larsen MR, Huganir RL. Experience-Induced Remodeling of the Hippocampal Post-synaptic Proteome and Phosphoproteome. Molecular & Cellular Proteomics. 2023;22.

66. Najar MA, Rex DAB, Modi PK, Agarwal N, Dagamajalu S, Karthikkeyan G, et al. A complete map of the Calcium/calmodulin-dependent protein kinase kinase 2 (CAMKK2) signaling pathway. J Cell Commun Signal. 2021;15:283–290.

67. Sheng M, Hoogenraad CC. The postsynaptic architecture of excitatory synapses: a more quantitative view. Annu Rev Biochem. 2007;76:823–847.

68. Uchinoumi H, Yang Y, Oda T, Li N, Alsina KM, Puglisi JL, et al. CaMKII-dependent phosphorylation of RyR2 promotes targetable pathological RyR2 conformational shift. J Mol Cell Cardiol. 2016;98:62–72.

69. Oh H, Lee S, Oh Y, Kim S, Kim YS, Yang Y, et al. Kv7/KCNQ potassium channels in cortical hyperexcitability and juvenile seizure-related death in Ank2-mutant mice. Nat Commun. 2023;14:3547.

70. Kim EE, Shekhar A, Lu J, Lin X, Liu F-Y, Zhang J, et al. PCP4 regulates Purkinje cell excitability and cardiac rhythmicity. J Clin Invest. 2014;124:5027–5036.

71. Alaimo A, Villarroel A. Calmodulin: A Multitasking Protein in Kv7.2 Potassium Channel Functions. Biomolecules. 2018;8:57.

72. Salvatore MF, Pruett BS, Dempsey C, Fields V. Comprehensive profiling of dopamine regulation in substantia nigra and ventral tegmental area. J Vis Exp. 2012:4171.

73. Lee AM, Mansuri MS, Wilson RS, Lam TT, Nairn AC, Picciotto MR. Sex Differences in the Ventral Tegmental Area and Nucleus Accumbens Proteome at Baseline and Following Nicotine Exposure. Front Mol Neurosci. 2021;14.

74. Smaga I, Jastrzębska J, Zaniewska M, Bystrowska B, Gawliński D, Faron-Górecka A, et al. Changes in the Brain Endocannabinoid System in Rat Models of Depression. Neurotox Res. 2017;31:421–435.

75. Qiu X-T, Guo C, Ma L-T, Li X-N, Zhang Q-Y, Huang F-S, et al. Transcriptomic and proteomic profiling of the anterior cingulate cortex in neuropathic pain model rats. Front Mol Neurosci. 2023;16:1164426.

76. Choi J-E, Lee J-J, Kang W, Kim HJ, Cho J-H, Han P-L, et al. Proteomic Analysis of Hippocampus in a Mouse Model of Depression Reveals Neuroprotective Function of Ubiquitin C-terminal Hydrolase L1 (UCH-L1) via Stress-induced Cysteine Oxidative Modifications. Mol Cell Proteomics. 2018;17:1803–1823.

77. Humo M, Lu H, Yalcin I. The molecular neurobiology of chronic pain–induced depression. Cell Tissue Res. 2019;377:21–43.

78. Shonesy BC, Bluett RJ, Ramikie TS, Báldi R, Hermanson DJ, Kingsley PJ, et al. Genetic Disruption of 2-Arachidonoylglycerol Synthesis Reveals a Key Role for Endocannabinoid Signaling in Anxiety Modulation. Cell Reports. 2014;9:1644–1653.

79. Fukumori *Ryo, Nakajima R, Ueo K, Yamaguchi T. ANXIOLYTIC AND ANTIDEPRESSANT EFFECTS OF INHIBITORS OF ENDOCANNABINOID DEGRADING ENZYME IN CHRONIC RESTRAINT-STRESSED MICE. International Journal of Neuropsychopharmacology. 2025;28:i325–i325.

80. Chen Z, Zheng L, Chen G. 2-Arachidonoylglycerol Attenuates Myocardial Fibrosis in Diabetic Mice Via the TGF-β1/Smad Pathway. Cardiovasc Drugs Ther. 2023;37:647–654.

81. Fang G, Wang Y. Effects of rTMS on Hippocampal Endocannabinoids and Depressive-like Behaviors in Adolescent Rats. Neurochem Res. 2018;43:1756–1765.

82. Brantl SA, Khandoga AL, Siess W. Mechanism of platelet activation induced by endocannabinoids in blood and plasma. Platelets. 2014;25:151–161.

83. Boczek T, Zylinska L. Receptor-Dependent and Independent Regulation of Voltage-Gated Ca2+ Channels and Ca2+-Permeable Channels by Endocannabinoids in the Brain. Int J Mol Sci. 2021;22:8168.

84. Zygmunt PM, Petersson J, Andersson DA, Chuang H, Sørgård M, Di Marzo V, et al. Vanilloid receptors on sensory nerves mediate the vasodilator action of anandamide. Nature. 1999;400:452–457.

85. Smart D, Gunthorpe MJ, Jerman JC, Nasir S, Gray J, Muir AI, et al. The endogenous lipid anandamide is a full agonist at the human vanilloid receptor (hVR1). Br J Pharmacol. 2000;129:227–230.

86. Karimi-Haghighi S, Shaygan M. Improvement in the Cognitive Function in Chronic Pain: Therapeutic Potential of the Endocannabinoid System. Mol Neurobiol. 2025. 10 March 2025. 10.1007/s12035-025-04814-8.

87. Blanton HL, Barnes RC, McHann MC, Bilbrey JA, Wilkerson JL, Guindon J. Sex differences and the endocannabinoid system in pain. Pharmacology Biochemistry and Behavior. 2021;202:173107.

88. Abd-Elsayed A, Jackson M, Gu SL, Fiala K, Gu J. Neuropathic pain and Kv7 voltage-gated potassium channels: The potential role of Kv7 activators in the treatment of neuropathic pain. Mol Pain. 2019;15:1744806919864256.

89. Aiba I, Noebels JL. Kcnq2/Kv7.2 controls the threshold and bi-hemispheric symmetry of cortical spreading depolarization. Brain. 2021;144:2863–2878.

90. Stewart AP, Gómez-Posada JC, McGeorge J, Rouhani MJ, Villarroel A, Murrell-Lagnado RD, et al. The Kv7.2/Kv7.3 heterotetramer assembles with a random subunit arrangement. J Biol Chem. 2012;287:11870–11877.

91. Vacher H, Trimmer JS. Trafficking mechanisms underlying neuronal voltage-gated ion channel localization at the axon initial segment. Epilepsia. 2012;53 Suppl 9:21–31.

92. Gomis-Perez C, Soldovieri MV, Malo C, Ambrosino P, Taglialatela M, Areso P, et al. Differential Regulation of PI(4,5)P2 Sensitivity of Kv7.2 and Kv7.3 Channels by Calmodulin. Front Mol Neurosci. 2017;10:117.

93. Hefting LL, D’Este E, Arvedsen E, Benned-Jensen T, Rasmussen HB. Multiple Domains in the Kv7.3 C-Terminus Can Regulate Localization to the Axon Initial Segment. Front Cell Neurosci. 2020;14:10.

94. Greene DL, Hoshi N. Modulation of Kv7 channels and excitability in the brain. Cellular and Molecular Life Sciences: CMLS. 2016;74:495.

95. Salzer I, Erdem FA, Chen W-Q, Heo S, Koenig X, Schicker KW, et al. Phosphorylation regulates the sensitivity of voltage-gated Kv7.2 channels towards phosphatidylinositol-4,5-bisphosphate. J Physiol. 2017;595:759–776.

96. van der Horst J, Greenwood IA, Jepps TA. Cyclic AMP-Dependent Regulation of Kv7 Voltage-Gated Potassium Channels. Front Physiol. 2020;11:727.

97. Workman ER, Niere F, Raab-Graham KF. Engaging homeostatic plasticity to treat depression. Mol Psychiatry. 2018;23:26–35.

98. Tien N-W, Kerschensteiner D. Homeostatic plasticity in neural development. Neural Dev. 2018;13:9.

99. Zolezzi DM, Larsen DB, McPhee M, Graven-Nielsen T. Effects of pain on cortical homeostatic plasticity in humans: a systematic review. Pain Rep. 2024;9:e1141.

100. Saneyoshi T, Wayman G, Fortin D, Davare M, Hoshi N, Nozaki N, et al. Activity-Dependent Synaptogenesis: Regulation by a CaM-Kinase Kinase/CaM-Kinase I/βPIX Signaling Complex. Neuron. 2008;57:94–107.

