## Supplementary Figure for "Molecular Dynamics in the Ventral Tegmental Area during Chronic Pain-Induced Negative Affect"

**Supplemental Figures**

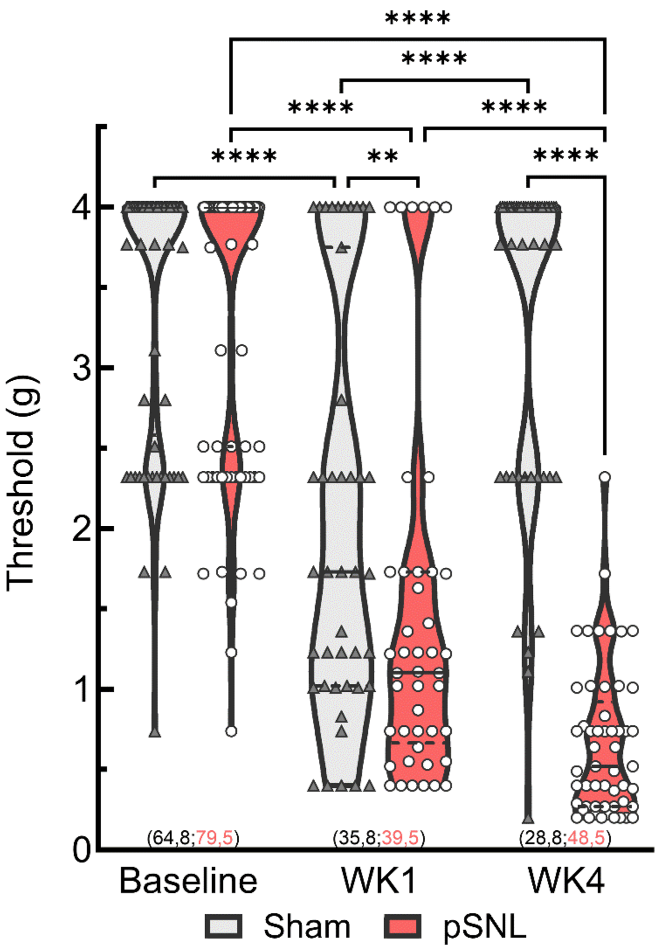


**Supplementary Figure 1. pSNL surgery causes long-lasting allodynia.** Violin plot depicts individual animal paw withdrawal threshold values in male and female mice from sham and pSNL-exposed groups (interaction $F_{(2,151)}$=39.46, ****$p$<0.0001). Both groups exhibited hypersensitivity at WK1 compared to baseline (****$p$<0.0001) to different extents (**$p$=0.0030). While pSNL mice continued to show hypersensitivity at WK4 (**$p$<0001), sham-exposed mice returned to withdrawal thresholds similar to baseline ****$p$<0.0001). Sample sizes are indicated in parentheses (Sham male, Sham female; pSNL male, pSNL female). Solid lines in violin plots depict median and dashed lines depict quartiles.

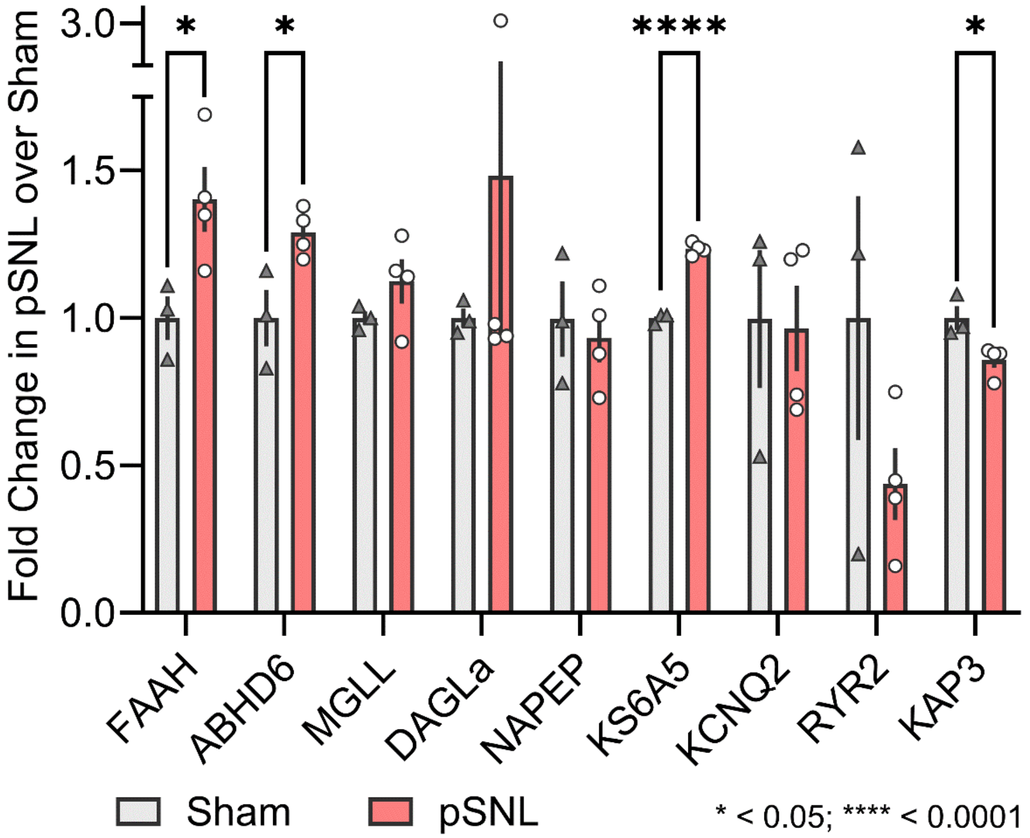


**Supplementary Figure 2. Early Imbalance in VTA Neuromodulation at WK1 pSNL.** Bar graph depicts the relative abundance of various proteins normalized to sham-operated mice (multiple unpaired t-tests). Included are enzymes involved in endocannabinoid (eCB) metabolism (fatty acid amide hydrolase (FAAH) and alpha/beta-hydrolase domain containing 6 (ABHD6)) and eCB synthesis (monoacylglycerol lipase (MGLL) and diacylglycerol lipase alpha (DAGLA)). Also analyzed are proteins important for anandamide production (N-acyl-phosphatidylethanolamine phospholipase D (NAPEPLD), and experience dependent synaptic plasticity (KS6KA5 (MSK1), and KAP3 (Prkar2b)). All data are expressed as mean ± SEM.

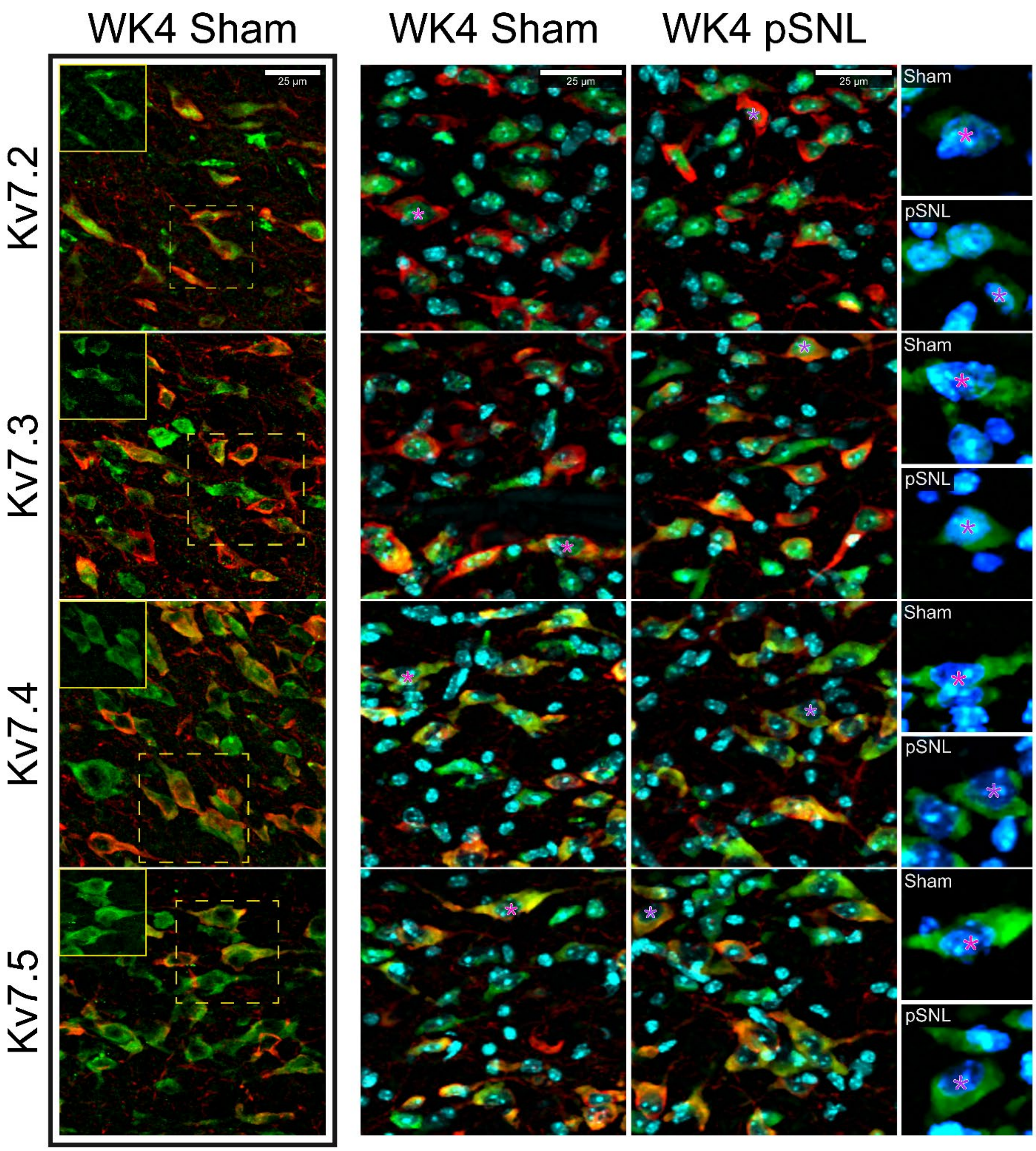


**Supplementary Figure 3. Cytosolic Expression Patterns of TH+ and Kv7.2-5 Among VTA cells.** High magnification confocal photomicrographs of the contralateral VTA region from male mice, divided into sham-operated (*left two panels*) and pSNL (*right panel*) groups, at four weeks post-surgery. Images depict co-localization of TH (*red*), Kv7.2-5 (*green*, presented in descending order), and nuclear DAPI staining (*blue*). Magnified images focusing on individual neurons (*far right*) show only Kv7 and DAPI staining. Images underwent rolling-ball background subtraction, linear contrast adjustment, and resizing for display. Scale bars represent 25 μm and asterisks represent the nucleus.
