## Supplementary Material 1 for "Molecular Dynamics in the Ventral Tegmental Area during Chronic Pain-Induced Negative Affect"

**Materials table**

| **REAGENT or RESOURCE** | **SOURCE** | **IDENTIFIER** |
| --- | --- | --- |
| Organism/Strain | | |
| Mouse: C57BL/6J | Jackson Labs (Waltham, MA, USA) | Strain #:000664  RRID: IMSR_JAX:000664 |
| Testing Compounds | | |
| 2-Arachidonylglycerol | Tocris Bioscience (Bristol, GBR) | Cat.# 1298 |
| Retigabine Dihydrochloride | Axon MedChem (Reston, VA, USA) | Cat.# 2252 |
| CS640 CaMKI Inhibitor | MedChemExpress (Monmouth Junction, NJ, USA) | Cat.# HY-148907 |
| IHC | | |
| *Antibodies and Counterstains* | | |
| Rabbit Polyclonal Anti-KCNQ2 | Alomone (Jerusalem, ISR) | Cat.# APC-050 |
| Rabbit Polyclonal Anti-KCNQ3 | Alomone (Jerusalem, ISR) | Cat.# APC-051 |
| Rabbit Polyclonal Anti-KCNQ4 | Alomone (Jerusalem, ISR) | Cat.# APC-164 |
| Rabbit Polyclonal Anti-KCNQ5 | Alomone (Jerusalem, ISR) | Cat.# APC-155 |
| Guinea Pig Polyclonal Anti-Tyrosine Hydroxylase | Synaptic Systems (Göttingen, DEU) | Cat.# 213-104 |
| Rabbit Monoclonal Anti-CaMKI | AbCam (Cambridge, GBR) | Cat.# ab68234 |
| Alexa Fluor 488 AffiniPure F(ab’)_2_ Fragment Donkey Anti-Rabbit IgG (H+L) | Jackson ImmunoResearch (West Grove, PA, USA) | Cat.# 711-546-152 |
| Cy3 AffiniPure Donkey Anti-Guinea Pig IgG (H+L) | Jackson ImmunoResearch (West Grove, PA, USA) | Cat.# 706-165-148 |
| DAPI (4',6-Diamidino-2-Phenylindole) | ThermoFisher (Waltham, MA, USA) | Cat.# D21490 |
| *Chemicals/Reagents* | | |
| Prolong Glass Antifade mountant | ThermoFisher (Waltham, MA, USA) | Cat.# P36980 |
| *Equipment* | | |
| SM2010R Microtome | Leica (Wetzlar, DEU) | N/A |
| Olympus FluoView FV1200 System | Olympus (Tokyo, JPN) | N/A |
| *Software* | | |
| Olympus FluoView FV10-ASW | Olympus (Tokyo, JPN) |  |
| FIJI/ImageJ (version 1.54p) | Schindelin et al. [1] | <https://imagej.net/software/fiji/> |
| Lipidomics | | |
| *Chemicals/Reagents* | | |
| d5-2-AG | Cayman Chemical (Ann Arbor, MI, USA) | Cat.# 362162 |
| d4-AEA | Cayman Chemical (Ann Arbor, MI, USA) | Cat.# 10011178 |
| *Equipment* | | |
| 6495C Triple Quadrupole Mass Spectrometer | Agilent (Palo Alto, CA, USA) | N/A |
| 1290 Infinity II UPLC System | Agilent (Palo Alto, CA, USA) | N/A |
| Acquity UPLC BEH C-18 1.7 µm 2.1 × 100 mm column | Waters (Milford, MA, USA) | SKU: 186002353 |
| Proteomics | | |
| *Chemicals/Reagents* | | |
| Digitonin | Sigma Aldrich (Burlington, MA, USA) | Cat.# D141 |
| Protease Inhibitor Cocktail | Sigma Aldrich (Burlington, MA, USA) | Cat.# P8340 |
| 100X Phosphatase inhibitor | Selleckchem (Houston, TX, USA) | Cat.# B15001 |
| *Kits* | | |
| Pierce BCA Protein Assay Kit | ThermoFisher (Waltham, MA, USA) | Cat.# 23225 |
| High Select Phosphopeptide Enrichment Kits & Reagents | Thermo Scientific (Waltham, MA, USA) | Cat. #: A32992 & A32993 |
| Pierce Quantitative Peptide Assays & Standards | Thermo Scientific (Waltham, MA, USA) | Cat. #: 23275 |
| *Equipment* | | |
| Orbitrap Fusion Lumos Tribrid Mass Spectrometer | Thermo Scientific (Waltham, MA, USA) | Cat.# IQLAAEGAAPFADBMBHQ |
| EASY-Spray Source | Thermo Scientific (Waltham, MA, USA) | Cat. #: ES081 |
| UltiMate 3000 RSLCnano System | Thermo Scientific (Waltham, MA, USA) | Cat. #: ULTIM3000RSLCNANO |
| EASY Spray C18 LC Column | Thermo Scientific (Waltham, MA, USA) | Cat. #: ES906 |
| Solid Phase C18 ZipTip | MilliporeSigma (Billerica, MA, USA) |  |
| *Software* | | |
| Progenesis QI for proteomics (version 2.4) | Nonlinear Dynamics Ltd. (Newcastle Upon Tyne, UK) | <https://www.nonlinear.com/progenesis/qi-for-proteomics/> |
| Mascot (version 2.6) | Matrix Science (London, UK) | <https://www.matrixscience.com/> |
| Perseus | Tyanova et al. [2]  Tyanova & Cox [3] | <https://maxquant.net/perseus/> |
| DAVID | Sherman et al. [4]  Huang et al. [5] | <https://davidbioinformatics.nih.gov/> |
| SynGO | Koopmans et al. [6] | <https://syngoportal.org/> |
| iceLogo | Colaert et al. [7] | <http://plogo.uconn.edu/> |
| Xcalibur (version 2.1.0) | Thermo Scientific (Waltham, MA, USA) | Cat. #: OPTON-30967 |
| Scaffold (version Scaffold_4.8.7) | Proteome Software (Portland, OR, USA) | <https://www.proteomesoftware.com/> |
| GraphPad Prism (version 10.4.2) | GraphPad Software (San Diego, California, USA) | <https://www.graphpad.com/> |

**Supplemental Methods**

**Animals**

A total of 154 mice (138 males and 16 females, Jackson Labs, C57BL/6J), aged 7-8 weeks, were kept in a climate-controlled room under a 12-hour light-dark cycle. They were housed in standard cages with three to five mice per cage and had ad libitum access to food and water. All procedures were approved by the University of Arizona Animal Care Use Committee and complied with NIH guidelines for laboratory animal care.

**Partial Sciatic Nerve Ligation (pSNL)**

Using published procedures [8], all 154 mice underwent surgery. For pSNL (n=80), mice were anesthetized with isoflurane then prepared for surgery. The sciatic nerve was exposed via a 2 mm cut made at the lateral aspect of the thigh followed by the separation of the muscle using extra-fine sharp tipped forceps. A surgical knot was tied around 1/3 of the width of the sciatic nerve using a 9-0 nylon suture. The muscle was then sutured using absorbable polyglycolic 5-0 suture then the skin using a non-absorbable polypropylene 6-0 suture. Sham surgeries (n=74) underwent a similar procedure; however, the nerve was not manipulated. Mice were administered gentamicin at a dose of 8 mg/kg (IP). The days following surgery mice were monitored for normal limb use. 3 mice were withdrawn from the study due to surgical complications.

**Mechanical Withdrawal Threshold**

We assessed tactile allodynia at baseline and at 1 week and 4 weeks post-surgery by measuring the paw withdrawal threshold in response to probing with a series of calibrated von Frey filaments, which serve as innocuous stimuli. Prior to testing, mice were acclimated in suspended wire-mesh cages for 60 minutes. von Frey filaments were then applied perpendicularly to the plantar surface of the paw for 2 seconds. A positive response was characterized by a sharp withdrawal of the paw during the initial application of the filaments (2.44, 2.83, 3.22, 3.61, 4.08, 4.31, 4.56). The paw withdrawal thresholds were determined using the non-parametric method described by Dixon (1980) [9], in which the stimulus intensity was incrementally increased until a positive response was observed, after which it was decreased until a negative response occurred. This protocol was repeated until three behavioral changes were noted using the “up and down” method [10, 11]. The 50% paw withdrawal threshold was determined as 10[Xf+kδ])/10,000, where Xf = the value of the last von Frey filament employed, k = Dixon value for the positive/negative pattern, and δ = the logarithmic difference between stimuli. To rapidly calculate 50% paw withdrawal thresholds, we used <https://bioapps.shinyapps.io/von_frey_app/>. Tactile allodynia was defined as a significant reduction in the paw withdrawal threshold compared to the pre-treatment baseline.

**Forced Swim Test**

Despair-like behavior was assessed 28 to 35 days post-surgery by measuring immobility time in a water-filled cylindrical container (25⁰C) for up to 5 minutes. Total immobility time throughout the duration of the test was used to evaluate despair-like behavior. One cohort received 2-AG (10 mg/kg, IP) shortly after preparation (1:1:8 DMSO:Tween 80:saline solution), 32 days post-surgery, and RTG (10 mg/kg, IP), prepared in 0.9% saline, 35 days post-surgery, administered 15 minutes before the FST. Another cohort received two doses of 2-AG (10 mg/kg, IP) 31 days post-surgery, both 15 and 60 minutes before the FST.

**Immunohistochemistry**

Mice were anesthetized using isoflurane, followed by cardiac perfusion with cold 0.1 M phosphate-buffered saline (PBS) and subsequently with cold 4% paraformaldehyde (PFA) in PBS solution. The brains were extracted and post-fixed in PFA for 24 hours, after which they were transferred to a 30% sucrose in 0.1 M PBS solution for an additional 48 hours. The midbrain was sectioned into coronal slices of 25 µm thickness using a SM2010R Leica microtome. Based on established criteria, the ventral tegmental area (VTA) was identified in the free-floating sections as the region medial to the compact part of the substantia nigra, while excluding the medial lemniscus (Franklin & Paxinos, 3rd Edition, 2007). Selected tissue slices underwent a washing procedure in 4% PFA for 20 minutes, followed by three sequential rinses in 0.1 M PBS and antigen retrieval conducted at 90°C for 60 minutes in 10 mM sodium citrate buffer containing 0.05% Tween-20 at pH 6.0. This was succeeded by two washes with PBS containing Triton X-100 (PBS-TX100) and one wash with PBS alone. The tissue was then incubated in a blocking solution composed of 3% bovine serum albumin (BSA), 0.3% Triton X-100, 0.3% Tween-20, 0.1% sodium azide (NaN₃), and 3% normal donkey serum (NDS) in 0.1 M PBS. Subsequently, VTA samples were placed in primary antibody solutions overnight with gentle oscillations at 4°C, including anti-KCNQ2 (1:800, Rabbit, Alomone), anti-KCNQ3 (1:800, Rabbit, Alomone), anti-KCNQ4 (1:400, Rabbit, Alomone), anti-KCNQ5 (1:200, Rabbit, Alomone), anti-CaMKIα (1:500, AbCam), and anti-tyrosine hydroxylase (TH) (1:800, Guinea Pig, Synaptic Systems). After the primary antibody incubation, the samples underwent PBS-TX100 washes then were incubated with secondary antibodies (anti-Guinea Pig Cy3, 1:600, Jackson ImmunoResearch; anti-Rabbit AF488, 1:400, Jackson ImmunoResearch) for one hour at room temperature. Counterstaining was performed using DAPI (ThermoFisher). Finally, the tissue sections were mounted onto glass slides using Prolong Glass Antifade mountant (ThermoFisher). The mounted samples were covered and allowed to harden overnight prior to confocal imaging.

**Imaging and Analysis**

Photomicrographs were acquired using Olympus FV10-ASW software (v4.02) on an Olympus Fluoview FV1200 confocal microscope set at 20× magnification (NA 0.8) and 1024 x 1024 pixel resolution. Throughout the imaging process, software and equipment settings were held constant for each antibody to ensure uniformity within each staining session. The ventral tegmental area (VTA) was delineated using coronal section schematic overlays sourced from the mouse brain atlas (Franklin & Paxinos, 3rd Edition, 2007). Subsequent image analyses were performed utilizing the FIJI/ImageJ software platform to determine the number of positive. A FIJI macro was created to quantify VTA cell counts for each protein. The macro included linear (Gaussian blur) and nonlinear (gamma correction, auto-threshold, rolling-ball background subtraction, and max intensity z-projection) adjustments to whole images to improve cell count accuracy and consistency. For each mouse specimen, quantifiable values for TH+, Kv7+, TH+/Kv7+, CaMKIα+, and TH+/CaMKIα+ were derived from 2-3 slices of the VTA that were contralateral to the surgical site.

**Quantification of 2-AG and AEA by LC-MS**

Following methodologies established in prior research [12], tissue samples (n = 4-5 bilateral punches per sample) were subjected to organic solvent extraction to purify for LC-MS analysis, following the protocol detailed by Wilkerson et al. [13]. On the day of processing, tissues were weighed and homogenized using a Dounce homogenizer with 1 ml of chloroform/methanol (2:1 v/v), supplemented with phenylmethylsulfonyl fluoride (PMSF, 1 mM) to inhibit degradation by endogenous enzymes. The homogenates were subsequently mixed with 0.3 ml of NaCl (0.7% w/v), vortexed, and centrifuged at 3,200 × g for 10 minutes at 4°C. The aqueous phase, along with debris, was collected and subjected to two additional extractions with 0.8 ml of chloroform. The organic phases were combined, and an internal standard was introduced to each sample. Mixed internal standards were prepared through serial dilution of d4-AEA and d5-2-AG in acetonitrile to facilitate concentration calculations and to account for run variability according to Wilkerson et al. [13].The organic solvents were evaporated using nitrogen gas, following which glycerol in methanol (6 µl, 30%) was added prior to evaporation. The dried samples were reconstituted with 0.2 ml of chloroform and combined with 1 ml of ice-cold acetone. The mixtures were then centrifuged at 1,800 × g for 5 minutes at 4°C, after which the organic layer of each sample was collected and subjected to further nitrogen evaporation.

Analysis of 2-AG and AEA was conducted using an Agilent 6495C triple quadrupole mass spectrometer coupled with a 1,290 Infinity II UPLC system (Agilent, Palo Alto, CA). The system was used in the electrospray positive mode with a gas temperature of 150°C, a flow rate of 5 L/min, a nebulizer pressure of 15 psi, a capillary voltage of 4,500 V, sheath gas set to 400°C at a flow rate of 12 L/min, and a nozzle voltage of 300 V. Monitored transitions included 348.3 → 287.3 and 62, 352.3 → 287.4 and 65.9, 379.3 → 287.2 and 269.2, and 384.3 → 287.2 and 296.1 for AEA, 2-AG, d4-AEA, and d5-2-AG, respectively. The first fragment was utilized for quantification, while the second served for confirmation. The initial 3 minutes of analysis were diverted to waste. Chromatographic separation was accomplished using an isocratic system comprising 21% 1 mM ammonium fluoride and 79% methanol on an Acquity UPLC BEH C-18 1.7 µm 2.1 × 100 mm column (Waters, Milford, MA) maintained at 60°C. Following each injection, the column was washed with 90% methanol for one minute and re-equilibrated for 5 minutes prior to the next injection. Samples were stored at 4°C. Calibration solutions were prepared from serial dilutions of AEA and 2-AG stock solutions in 80% C₂H₃N. Calibration curves for each analysis were generated by adding 10 µl of internal standard solution to 20 µl of standard solution. To the dried samples, 200 µl of a solvent mixture containing 80:20 C₂H₃N:H₂O was added, followed by vortexing and sonication. Samples underwent centrifugation at 15,800 × g for 5 minutes at 4°C, whereupon the supernatant was transferred to autosampler vials, and 5 µl was injected for analysis.

**In-gel digestion**

Using previously published procedures [14–16], pooled VTA samples (2-3 bilateral punches/sample) from 1 and 4 weeks post-surgery were collected on ice, snap froze on dry ice, and stored at -80⁰C. Radioimmunoprecipitation assay buffer (RIPA; 20 mM Tris HCL, 150 mM NaCl, 2 mM EDTA, 0.1% SDS, 1% TritonX-100, 0.25% Deoxycholate, 1 mM Na Orthovanadate, 1 mM phenylmethylsulfonyl fluoride) with 1x protease inhibitor (SigmaAldrich), 1x phosphatase inhibitor (Selleckchem), and 1% Digitonin (SigmaAldrich) was added to pooled samples (2-3 bilateral punches/sample) and iced for 15 min. Samples were homogenized by ultrasonication and centrifuged 4⁰C at 15000g for 10 minutes. Protein concentration in the supernatant was determined using the Pierce BCA protein assay kit (ThermoFisher).

For the proteome wide experiments, 50 μg of mouse VTA lysate was separated by SDS-PAGE, and each lane was cut into seven slices. The gel slices were subjected to trypsin digestion and the resulting peptides were purified by C^18^-based desalting exactly as previously described [14, 15]. In brief, the SDS-PAGE gel slices were placed in a 0.6 mL LoBind polypropylene tube (Eppendorf), destained twice with 375 μL of 50% acetonitrile (ACN) in 40 mM NH_4_HCO_3_ and dehydrated with 100% acetonitrile (ACN) for 15 min. After removal of the ACN by aspiration, the gel pieces were dried in a vacuum centrifuge at 60°C for 30 minutes. Trypsin (250 ng; Sigma-Aldrich) in 20 μL of 40 mM NH_4_HCO_3_ was added, and the samples were maintained at 4°C for 15 minutes prior to the addition of 50-100 μL of 40 mM NH_4_HCO_3_. The digestion was allowed to proceed at 37°C overnight and was terminated by addition of 10 mL of 5% formic acid (FA). After further incubation at 37°C for 30 minutes and centrifugation for 1 minute, each supernatant was transferred to a clean LoBind polypropylene tube. The extraction procedure was repeated using 40 mL of 0.5% FA, and the two extracts were combined and dried down to ∼5-10 mL followed by the addition of 10 mL of 0.05% heptafluorobutyric acid/5% FA (vol/vol) and incubation at room temperature for 15 minutes. The resulting peptide mixtures were loaded on a solid phase C18 ZipTip (MilliporeSigma, Billerica, MA) and washed with 35 mL 0.005% heptafluorobutyric acid/5% FA (vol/vol) followed by elution first with 4 mL of 50% ACN/1% FA (vol/vol) and then a more stringent elution with 4 mL of 80% ACN/1% FA (vol/vol). The eluates were combined and dried completely by vacuum centrifugation and 6 mL of 0.1% FA (vol/vol) was added followed by sonication for 2 minutes. 2.5 mL of the final sample was then analyzed by mass spectrometry.

**Mass spectrometry**

HPLC-ESI-MS/MS was performed in positive ion mode on a Thermo Scientific Orbitrap Fusion Lumos tribrid mass spectrometer fitted with an EASY-Spray Source (Thermo Scientific) as previously described [15]. In brief, NanoLC was performed using a Thermo Scientific UltiMate 3000 RSLCnano System with an EASY Spray C18 LC column (Thermo Scientific, 50 cm x 75 mm inner diameter, packed with PepMap RSLC C18 material, 2 mm, cat. # ES803); loading phase for 15 minutes; mobile phase, linear gradient of 1–47% ACN in 0.1% FA for 106 minutes, followed by a step to 95% ACN in 0.1% FA over 5 minutes, hold 10 minutes, and then a step to 1% ACN in 0.1% FA over 1 minute and a final hold for 19 minutes (total run 156 minutes); Buffer A = 100% H_2_O in 0.1% FA; Buffer B = 80% ACN in 0.1% FA; flow rate, 300 nL/min. All solvents were liquid chromatography mass spectrometry grade. Spectra were acquired using Xcalibur, version 2.1.0 (Thermo Scientific). A “TopSpeed” data-dependent MS/MS analysis was performed (acquisition of a full scan spectrum followed by collision-induced dissociation mass spectra of the Top N most intense precursor ions within the 3 second cycle time). Dynamic exclusion was enabled with a repeat count of 1, a repeat duration of 30 seconds, an exclusion list size of 500, and an exclusion duration of 40 seconds.

**Label-free Quantitative Proteomics**

Progenesis QI for proteomics software (version 2.4, Nonlinear Dynamics Ltd., Newcastle upon Tyne, UK) was used to perform ion-intensity based label-free quantification similar to as previously described [15]. In brief, in an automated format, .raw files were imported and converted into two-dimensional maps (y-axis = time, x-axis =m/z) followed by selection of a reference run for alignment purposes. An aggregate data set containing all peak information from all samples was created from the aligned runs, which was then further narrowed down by selecting only +2, +3, and +4 charged ions for further analysis. A peak list of fragment ion spectra was exported in Mascot generic file (.mgf) format and searched against the *Mus musculus* SwissProt database (17079 entries) using Mascot (Matrix Science, London, UK; version 2.6). The search variables that were used were: 10 ppm mass tolerance for precursor ion masses and 0.5 Da for product ion masses; digestion with trypsin; a maximum of two missed tryptic cleavages; variable modifications of oxidation of methionine and phosphorylation of serine, threonine, and tyrosine; ^13^C=1. The resulting Mascot .xml file was then imported into Progenesis, allowing for peptide/protein assignment, while peptides with a Mascot Ion Score of <25 were not considered for further analysis. Precursor ion-abundance values for peptide ions were normalized to all proteins. Unbiased hierarchical clustering analysis (heat map) and principal component analysis (PCA) were conducted using Perseus [2, 3]. Gene ontology (GO) and Reactome pathway enrichment analyses were completed with the DAVID tool [5]. For synapse-specific cellular component GO analysis, SynGO was utilized. Phosphorylation sequence enrichment analysis was performed using the iceLogo generation tool [7]. Volcano plots and scatter plots were created in GraphPad Prism.

**Phosphoproteomics**

To evaluate differences in protein phosphorylation abundance with pSNL and CS640 treatment, mice were anesthetized and administered a 5 μL intracerebroventricular injection of CS640 (2 μg/μL) or vehicle (1:1:8 DMSO:Tween 80:saline solution). The researchers who administered the drug and vehicle were blinded to group assignments. Bilateral VTAs were then collected 30 minutes later. 0.2-0.4 mg of VTA protein lysate per sample underwent in-solution tryptic digestion and phosphopeptide enrichment using metal oxide affinity chromatography per the manufacturer’s protocol (Thermo Scientific) similar to as previously described [17, 18]. The dried peptides were resuspended in 20 μL of 0.1% FA (v/v) and the peptide concentration was determined with the Pierce Quantitative Colorimetric Peptide Assay Kit per the manufacturer’s protocol (Thermo Scientific). 350 ng of the final sample was then analyzed by mass spectrometry.

**Statistical Analysis**

Statistical analyses were conducted using GraphPad Prism 10.4.2, employing two- and three-way ANOVAs (mixed-effects (ME) where appropriate), with Tukey's or Fisher's LSD post hoc tests for group comparisons and unpaired t-tests where appropriate. All GraphPad Prism statistics are presented in **Supplementary Table 1**.

**References**

1.	Schindelin J, Arganda-Carreras I, Frise E, Kaynig V, Longair M, Pietzsch T, et al. Fiji: an open-source platform for biological-image analysis. Nat Methods. 2012;9:676–682.

2.	Tyanova S, Temu T, Sinitcyn P, Carlson A, Hein MY, Geiger T, et al. The Perseus computational platform for comprehensive analysis of (prote)omics data. Nat Methods. 2016;13:731–740.

3.	Tyanova S, Cox J. Perseus: A Bioinformatics Platform for Integrative Analysis of Proteomics Data in Cancer Research. Methods Mol Biol. 2018;1711:133–148.

4.	Sherman BT, Hao M, Qiu J, Jiao X, Baseler MW, Lane HC, et al. DAVID: a web server for functional enrichment analysis and functional annotation of gene lists (2021 update). Nucleic Acids Res. 2022;50:W216–W221.

5.	Huang DW, Sherman BT, Lempicki RA. Systematic and integrative analysis of large gene lists using DAVID bioinformatics resources. Nat Protoc. 2009;4:44–57.

6.	Koopmans F, van Nierop P, Andres-Alonso M, Byrnes A, Cijsouw T, Coba MP, et al. SynGO: An Evidence-Based, Expert-Curated Knowledge Base for the Synapse. Neuron. 2019;103:217-234.e4.

7.	Colaert N, Helsens K, Martens L, Vandekerckhove J, Gevaert K. Improved visualization of protein consensus sequences by iceLogo. Nat Methods. 2009;6:786–787.

8.	Korah HE, Cheng K, Washington SM, Flowers ME, Stratton HJ, Patwardhan A, et al. Partial Sciatic Nerve Ligation: A Mouse Model of Chronic Neuropathic Pain to Study the Antinociceptive Effect of Novel Therapies. J Vis Exp. 2022. 6 October 2022. https://doi.org/10.3791/64555.

9.	Dixon WJ. Efficient analysis of experimental observations. Annu Rev Pharmacol Toxicol. 1980;20:441–462.

10.	Chaplan SR, Bach FW, Pogrel JW, Chung JM, Yaksh TL. Quantitative assessment of tactile allodynia in the rat paw. J Neurosci Methods. 1994;53:55–63.

11.	Chaplan SR, Pogrel JW, Yaksh TL. Role of voltage-dependent calcium channel subtypes in experimental tactile allodynia. J Pharmacol Exp Ther. 1994;269:1117–1123.

12.	Liktor-Busa E, Levine AA, Palomino SM, Singh S, Wahl J, Vanderah TW, et al. ABHD6 and MAGL control 2-AG levels in the PAG and allodynia in a CSD-induced periorbital model of headache. Front Pain Res (Lausanne). 2023;4:1171188.

13.	Wilkerson JL, Niphakis MJ, Grim TW, Mustafa MA, Abdullah RA, Poklis JL, et al. The Selective Monoacylglycerol Lipase Inhibitor MJN110 Produces Opioid-Sparing Effects in a Mouse Neuropathic Pain Model. J Pharmacol Exp Ther. 2016;357:145–156.

14.	Kruse R, Krantz J, Barker N, Coletta RL, Rafikov R, Luo M, et al. Characterization of the CLASP2 Protein Interaction Network Identifies SOGA1 as a Microtubule-Associated Protein. Mol Cell Proteomics. 2017;16:1718–1735.

15.	Parker SS, Krantz J, Kwak E-A, Barker NK, Deer CG, Lee NY, et al. Insulin Induces Microtubule Stabilization and Regulates the Microtubule Plus-end Tracking Protein Network in Adipocytes. Mol Cell Proteomics. 2019;18:1363–1381.

16.	Vizcarra VS, Barber KR, Franca-Solomon G, Majuta L, Smith A, Langlais PR, et al. Targeting 5-HT2A receptors and Kv7 channels in PFC to attenuate chronic neuropathic pain in rats using a spared nerve injury model. Neurosci Lett. 2022;789:136864.

17.	Keresztes A, Olson K, Nguyen P, Lopez-Pier MA, Hecksel R, Barker NK, et al. Antagonism of the mu-delta opioid receptor heterodimer enhances opioid antinociception by activating Src and calcium/calmodulin-dependent protein kinase II signaling. Pain. 2022;163:146–158.

18.	Levine AA, Liktor-Busa E, Balasubramanian S, Palomino SM, Burtman AM, Couture SA, et al. Depletion of Endothelial-Derived 2-AG Reduces Blood-Endothelial Barrier Integrity via Alteration of VE-Cadherin and the Phospho-Proteome. Int J Mol Sci. 2023;25:531.
